# Oligonucleotide-based CRISPR-Cas9 toolbox for efficient engineering of *Komagataella phaffii*

**DOI:** 10.1101/2023.12.09.570913

**Authors:** Tomas Strucko, Adrian-E Gadar-Lopez, Frederik B Frøhling, Emma T Frost, Helen Olsson, Zofia D Jarczynska, Uffe H Mortensen

## Abstract

*Komagataella phaffii* (*Pichia pastoris*) is a methylotrophic yeast that is favored by industry and academia mainly for expression of heterologous proteins. However, its full potential as a host for bio-production of valuable compounds is not yet fully exploited. The emergence of CRISPR-Cas9 technology has significantly improved the efficiency of gene manipulations of non-conventional species including *K. phaffii*. Yet, improvements in gene-editing methods are desirable to further accelerate engineering of industrially and scientifically relevant *K. phaffii* strains. In this study, we have developed a versatile one vector-based CRISPR-Cas9 method and showed that it works efficiently at different genetic loci using linear DNA fragments with very short targeting sequences. Importantly, we show that by using our setup it is possible to catalyze single-stranded oligonucleotide-mediated mutagenesis and marker-free gene integrations. Notably, we performed site-specific point mutations and full gene deletions using single stranded 90-mers at very high efficiencies. Lastly, we present a strategy for transient inactivation of non-homologous end-joining (NHEJ) pathway, where *KU70* gene is disrupted by a visual marker (*uidA* gene). The latter system enables precise CRISPR-Cas9 based editing (including multiplexing) and accelerates selection of the mutants that have simultaneously undergone a desired genetic modification(s) and restored NHEJ-proficient genotype. In conclusion, the tools presented in this study can be applied for easy and efficient engineering of *K. phaffii* strains and could potentially be coupled with high-throughput automated workflows.

## Introduction

*Komagataella phaffii* (aka. *Pichia pastoris*) is a methylotrophic yeast that has gained its popularity as a preferred host for heterologous protein expression and to this date is being widely used for academic and industrial purposes (Werten et al., 2019; Yang and Zhang, 2018). In recent years, *K. phaffii* became increasingly applied as a cell factory for production of small value-added molecules using various carbon sources as substrates reviewed in (Peña et al., 2018; Yang and Zhang, 2018). More importantly, the use of one carbon substrate (such as methanol) for bioconversion or production of high value chemicals was successfully implemented. For example, Gassler et al (Gassler et al., 2020) has recently demonstrated that *K. phaffii* can be successfully engineered to assimilate CO_2_. In addition, relevant physiological characteristics have boosted *K. phaffii* application in fundamental research (Bernauer et al., 2021; Karbalaei et al., 2020). Thus, with rapidly growing interest in this organism, the development of efficient methods for gene editing that are compatible with high throughput experiments are necessary.

Gene editing in the *K. phaffii* is hampered by the dominance of the non-homologous end joining (NHEJ) DNA repair pathway, which favors random integration over homologous recombination mediated gene targeting (Näätsaari et al., 2012). The emergence of CRISPR-Cas9 technology has revolutionized gene engineering of many non-model yeasts (Cai et al., 2019; Stovicek et al., 2017) and has also been successfully adapted to *K. phaffii* (Weninger et al., 2016) where the repertoire of CRISPR-Cas9 tools steadily increases (Dalvie et al., 2020; Gu et al., 2019; Liao et al., 2021; Liu et al., 2019; Wang et al., 2023). DNA double strand breaks induced by a CRISPR nuclease allows easy gene inactivation via erroneous NHEJ based repair, as well as efficient template guided gene editing mediated by HR. However, recent studies have demonstrated that CRISPR nuclease induced DNA double strand breaks, despite the intended edits, also induce unwanted off-target mutations and/or result in gross chromosomal rearrangements in NHEJ-proficient strain backgrounds (Garrigues et al., 2023; Schusterbauer et al., 2022). Thus to minimize occurrence of unwanted mutations and improve gene targeting efficiency it is desirable to inactivate the NHEJ pathway. However, elimination of NHEJ comes at a cost of reduced transformation- and growth rates as well as increased sensitivity to radiation and other DNA-damaging conditions (Carvalho et al., 2010; Näätsaari et al., 2012). After CRISPR based cell factory construction in an NHEJ deficient background, it is therefore advisable to restore the NHEJ pathway prior to thorough performance evaluation. Lastly, improved methods for construction of gene targeting substrates (GTS) that are necessary for large scale metabolic engineering experiments in *K. phaffii* are needed.

In this work, we have developed a highly efficient CRISPR-Cas9 method for engineering of prototrophic *K. phaffii* strains. Our method is based on one plasmid system that facilitates simultaneous co-expression of Cas9 and single or multiple arrays of sgRNA cassettes. We have evaluated our method by modifying multiple genetic loci with linear DNA fragments equipped with homology arms of varying size. Importantly, we have demonstrated that short (90nt) single-stranded DNA oligonucleotides are sufficient to mediate repair of Cas9 induced double-strand breaks (DSBs), thus significantly reducing time and cost needed for construction of complex GTSs. Moreover, we present a method for transient disruption of the NHEJ pathway by inserting the visually selectable *uidA* marker into *KU70*. The system enables precise Cas9-mediated gene modifications and simultaneous, and easy to detect, recovery of the NHEJ pathway to the wild type state. Moreover, the color marker associated with the NHEJ pathway facilitates effortless selection of yeast colonies that are proficient for homologous recombination, and which have therefore most likely undergone the intended genetic manipulations.

## Materials and Methods

### Strains and cultivation media

*Escherichia coli* DH5α strain was used for propagation and assembly of plasmids. In all cases, *E. coli* strains were were cultivated in Lysogeny Broth (Bertani, 1951) medium supplemented with 100 mg/L of ampicillin (Sigma).

All yeast experiments were done in a wild-type strain of *Komagataella phaffii* (CBS 2612) and its NHEJ-deficient derivatives sDIV390 (*ku70* point mutation, this study) and sDIV291 (*ku70::uidA*, this study). For propagation and transformation of *K. phaffii*, a yeast extract peptone dextrose (YPD) medium containing 1% yeast extract, 2% peptone and 2% glucose (and 2% of agar for solid media) was used. For the selection of transformed yeast strains, YPD medium was supplemented with 100 mg/L of nourseothricin (ClonNAT; Werner BioAgents). For blue colony screen (*uidA* marker), solid medium was additionally supplemented with 60 mg/L X-Gluc (Thermo Fisher Scientific).

### Molecular cloning techniques

All DNA fragments necessary for plasmid or gene targeting substrate (GTS) construction were amplified by PCR using Phusion U Master Mix (ThermoFisher) according to the manufacturer’s recommendations. All primer sequences used in this study are listed in Supplementary Table S1 and were purchased from Integrated DNA Technologies (IDT). All PCR fragments were gel-purified using NucleoSpin Gel and PCR Clean-up Kit (MACHEREY-NAGEL) by following manufacturer’s protocol. New plasmids used in this work were constructed by the uracil-specific excision reagent (USER™) by following well established protocols (Nour-Eldin et al., 2006). Plasmid minipreps were prepared using GeneElute Plasmid MiniPrep Kit (Sigma-Aldrich) according to the manufacturer’s protocol. New DNA constructs were validated by Sanger sequencing using the Mix2Seq service (Eurofins). All plasmids used and constructed in this work are listed in Supplementary Table S2. Synthetic DNA sequences were ordered as gBlocks from IDT (https://eu.idtdna.com).

### DNA assembly methods

#### Construction of sgRNA and Cas9 expressing plasmids

The basic Cas9 expression plasmid (pDIV151) was assembled by cloning *GAP1* promoter amplified from genomic DNA (gDNA) of *K. phaffii* (CBS 2612), and *cas9-ScCYC1t* cassette amplified from plasmid pCfB2312 (Stovicek et al., 2015), into AsiSI/Nb.bsmI linearized vector pDIV019 (Strucko et al., 2021). The basic sgRNA-Cas9 plasmid (pDIV153) was constructed by inserting three fragments: the *TEF1* promoter amplified from *K. phaffii* gDNA, a s*gRNA-tRNA* construct amplified from pFC902 plasmid (Nødvig et al., 2018), and the *AOX* terminator from *K. phaffii* gDNA, into AsiSI/Nb.bsmI digested pDIV151 vector. Each sgRNA-Cas9 vector expressing a single target specific guide sequence was assembled as follows: two PCRs fragments amplified from the pDIV153 plasmid were USER-cloned into the linearized pDIV151 backbone, resulting in the following plasmids pDIV259 (sgRNA1-IS1), pDIV260 (sgRNA1-IS3), pDIV270 (sgRNA1-KU70), pDIV272 (sgRNA2-uidA), pDIV273 (sgRNA1-ADE2) and pDIV274 (sgRNA2-ADE2). A Cas9 vector expressing two gRNAs (*PEP4* and *uidA*) was assembled by cloning three PCRs fragments amplified from pDIV153, resulting in a pDIV602 plasmid. A detailed cloning procedure for all Cas9 and sgRNA expressing vectors is depicted in the Supplementary Figure S1. The relevant plasmids and their maps will be available at https://www.addgene.org/Uffe_Mortensen/upon publication of this manuscript.

#### Construction of plasmids harboring gene targeting substrates (GTS)

The assembly of *mVenus* expressing cassettes flanked by long homology (LH) sequences guiding to specific loci in the genome of *K. phaffii* was done in two steps. First, an mVenus expressing plasmid (pDIV479) was constructed by cloning *TEF1* promoter amplified from *K. phaffii* gDNA, and an *mVenus-ScCYC1t* cassette amplified from a synthetic gBlock (IDT), into a PacI/Nt.bbvCI linearized vector pAC125 (Hansen et al., 2011). Second, locus specific upstream and downstream homology arms were amplified from *K. phaffii* gDNA, and individually USER-cloned into a PacI/Nt.bbvCI linearized pDIV479 vector. The two resulting plasmids were denoted as pDIV505 (for integration site IS1), and pDIV506 (for integration site IS3). Similarly, the assembly of the *uidA* expressing cassette for disruption of *KU70* homolog was done in two steps. First, an uidA expressing plasmid (pDIV033) was constructed by USER-cloning *TEF1* promoter amplified from *S. cerevisiae* gDNA, and an *uidA-ScCYC1t* cassette amplified from a synthetic gBlock (IDT), into AsiSI/Nb.bsmI linearized pDIV019 vector. Second, the *KU70* disrupting vector (pDIV643) was constructed by cloning *KU70* locus-specific upstream and downstream homology sequences amplified from *K. phaffii* gDNA, and *ScTEF1p-uidA-CYC1t* cassette, amplified from pDIV033, into PacI/Nt.bbvCI linearized pAC125 vector. A detailed cloning procedure for all GTS expressing vectors is depicted in the Supplementary Figure S2. Prior to transformation into *K. phaffii*, plasmids pDIV505, pDIV506 and pDIV643 were linearized with NotI enzyme in order to liberate linear DNA fragments (GTSs).

#### Construction of PCR based GTS

The *mVenus* expressing cassettes flanked by short homology (SH) sequences guiding to specific loci in the genome of *K. phaffii* were PCR-amplified (template pDIV479) by using primers with 60bp tails homologous to upstream and downstream regions of IS1 and IS3 loci. A GTS used to restore the wild-type KU70 genotype was PCR-amplified from *K*.*phaffii* gDNA with a specific primer pair PR_DIV2660 and PR_DIV2661. A detailed procedure for construction of PCR-based GTSs is depicted in the Supplementary Figure S3.

### Design of 20 nt guiding sequences of sgRNAs

For each genetic modification, a specific 20-nt sequence (see Table 1) guiding the Cas9 nuclease to the appropriate genetic locus was designed using CRISPRdirect (https://crispr.dbcls.jp/); an online tool developed by Naito et al. (Naito et al., 2015).

**Table 1.**
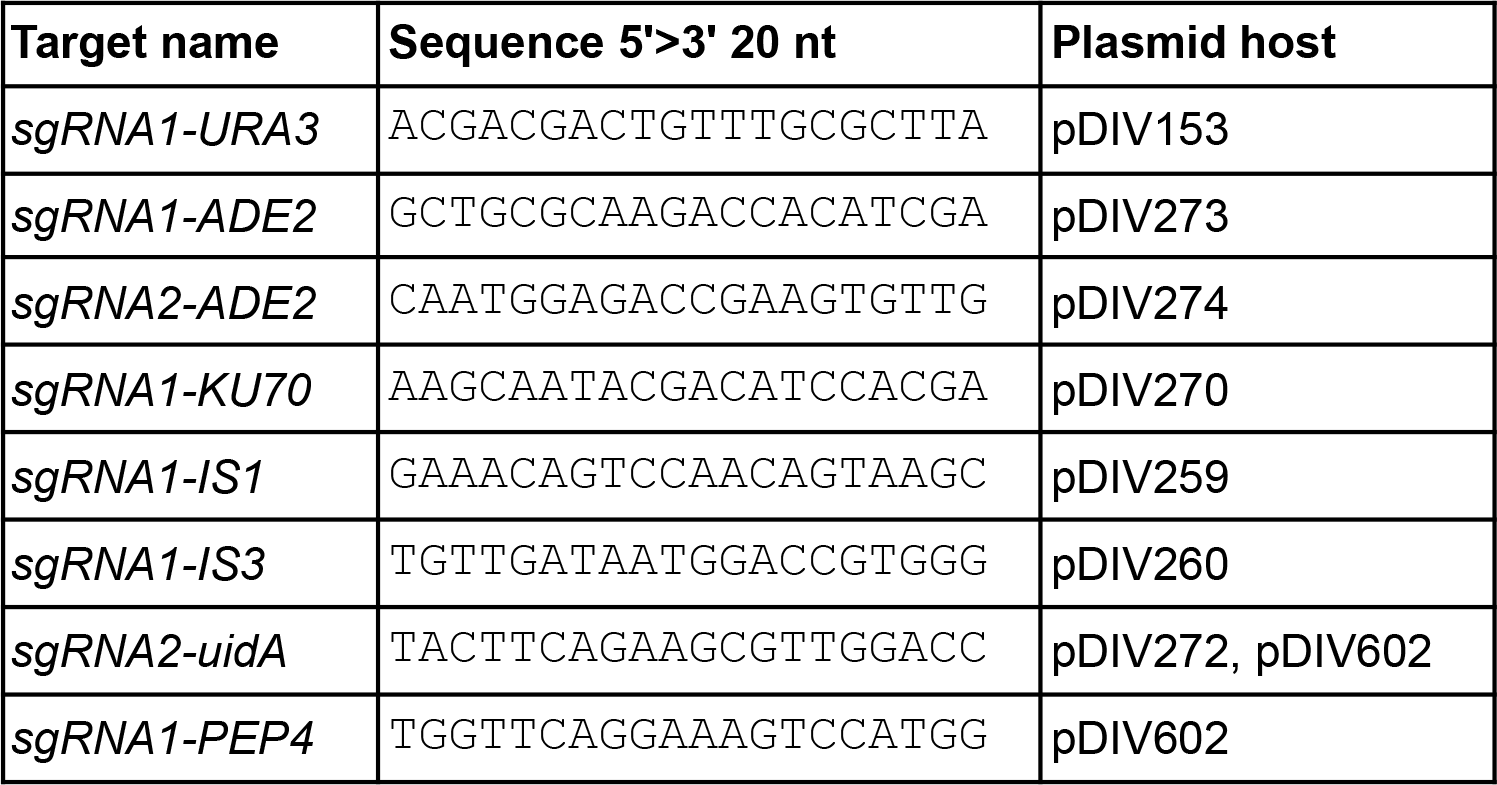
Targeting sgRNA sequences used in this study.

### Transformation of *K. phaffii*

All *K. phaffii* strains were transformed using an electroporation method slightly modified from (Lin-Cereghino et al., 2005). In brief, a single colony was inoculated into 5 mL of YPD media and incubated O/N at 30°C with 200 rpm shaking. Next day, appropriate volume of O/N culture was transferred into a fresh 50 mL of YPD in 500 mL baffled shake flask to a starting OD of ∼0.1. Fresh cultures were incubated at 30°C with orbital shaking at 200 rpm until OD of ∼1.5 was achieved (approx. 5 hours). Cells were collected by centrifugation at 3000 xg for 5 minutes, the supernatant was discarded and the pellet was resuspended in 20 mL of Transformation Buffer (100 mM LiAc, 10 mM DTT, 0.6 M sorbitol, and 6 mM Tris HCl pH 8.5) and incubated at room temperature for 30 minutes. Cells were centrifuged at 3000 xg for 5 minutes, and the supernatant was discarded. Pellet was resuspended in 1.5 mL 1M ice-cold sorbitol and transferred to sterile 2mL Eppendorf tubes. The pellet was washed three times by centrifuging them at 10000 xg for 30 seconds and then resuspended in 2 mL of 1M ice-cold sorbitol. The cell suspension was distributed into 10 sterile 1.5mL Eppendorf tubes (200 uL per vial) and used for transformation, or frozen at -80°C for later use. Routinely, ∼0.2 pmol of plasmid was added to the mix, and 1 pmol of linear DNA was added when required. The cell-DNA mix was incubated on ice for 5 minutes and transferred to an ice-cold mm electroporation cuvette. Cells were electroporated by Bio-Rad MicroPulser Electroporator using predefined “Pic” settings for *Pichia pastoris* (1 pulse at 2 kV) and immediately transferred into 2 mL Eppendorf tubes containing 0.8 mL of ice-cold YPD (containing 1M Sorbitol). The cell suspension was incubated for 3 hours at 30°C without shaking and plated on selective media. Transformation plates were incubated for 3-5 days at 30°C. Each transformation experiment was done in four replicates.

### *K. phaffii* mutant strain construction

The two NHEJ deficient *K. phaffii* strains were constructed as follows: first, a wild-type strain sDIV0084 (CBS 2612) was transformed with a pDIV270 plasmid and incubated on YPD+NTC media. The resulting transformants were streak-purified in YPD+NTC media, and the *KU70* locus of eight random colonies was Sanger sequenced (Supplementary Figure S4). One colony that contained a single nucleotide deletion (A32Δ) in *KU70* gene was saved and named as sDIV0360. Second, a wild-type strain sDIV0084 was co-transformed with a pDIV270 plasmid and linear uidA expressing GTS (NotI linearized and gel purified from pDIV643 plasmid), and incubated on YPD+NTC+X-Gluc media. Four blue colonies were streak-purified on YPD+NTC+X-Gluc medium, and correct GTS integration was confirmed by colony PCR (see text below). One validated colony (*ku70::uidA*) was saved and named sDIV291. Strains sDIV291 and sDIV390 were cured for sgRNA-Cas9 plasmids prior to their application for upcoming experiments.

### Validation of *K. phaffii* mutant strains

*K. phaffii* transformants were screened for expected phenotypes (colony color or fluorescence) on selection plates. In the case of *ade2*, plates were visually inspected by counting the number of red colonies with regard to the total number of colonies formed. For the selection of strains expressing the *uidA* gene, transformants were plated on solid media supplemented with X-Gluc and formation of blue/white colonies were assessed. In the case of integration of the *mVenus* gene coding for yellow fluorescence protein (YFP), transformation plates were photographed using a Light and Filter set (NIGHTSEA™) optimized for yellow fluorophores. Subsequently, images were evaluated by counting the number of fluorescent colonies with regard to the total number of colonies formed. Mutant strains showing the correct phenotype were tested by colony PCR to confirm introduction of the desired mutation(s). Routinely, eight random colonies per each biological replicate were selected for PCR analysis. All colony PCRs were done using Quick-Load® Taq 2X Master Mix (New England Biolabs) according to the manufacturer’s recommendation. Prior running the PCR reaction cells were thermally pretreated (98°C for 5 min, 4°C for 30 sec, 98°C for 5 min, and 80°C for 10 min). Sizes of DNA fragments were analyzed by running 1% agarose gel electrophoresis. In cases where DNA sequence needed to be determined, a PCR fragment of interest was PCR column-purified and sent for Sanger Sequencing using Mix2Seq service (Eurofins).

### Agar plate image acquisition

Images of agar plates with *K. phaffii* colonies were photographed using an SLR camera (Nikon D90). Settings for the white light photography were as follows: ISO Speed - ISO250, F-stop - f/7.1, Exposure time -1/80 s, Focal length - 60 mm. Yellow (mVenus) fluorescence was examined using an in-house built digital camera setup with a Light and Filter set (NIGHTSEA™), for details refer to (Vanegas et al., 2019). For acquisition of fluorescent images in all cases the exposure time was fixed to 1.6 s and the remaining settings were as follows: ISO Speed - ISO200, F-stop - f/7.1, Focal length - 60 mm.

## Results and Discussion

### CRISPR-Cas9 tool design and validation

The strength of the CRISPR-Cas technology is that its sgRNA guided CRISPR endonuclease can be effortlessly re-programmed to cleave at almost any sequence in the genome to set the stage for targeted gene editing. However, the success and efficiency of CRISPR tools relies highly on well-balanced expression of specific sgRNA and Cas9, and often requires multiple steps of optimization as reported in the first example for *K. phaffii* (Weninger et al., 2016). In addition, features (i.e. ARS, backbone and selection marker) of a vector that facilitates the CRISPR-Cas9 expression plays a key role in the success of editing (Gu et al., 2019). Thus, based on the accumulated knowledge, we have designed a plasmid-based CRISPR-Cas9 gene-editing system for *K. phaffii* (Figure 1**a** and Materials and Methods). Specifically, we made a CRISPR vector, based on pDIV019 backbone, which is able to replicate in a selectable manner in *K. phaffii* as it contains a synthetic autonomously replicating sequence (h-ARS) and a dominant NatMX marker, which are both compatible with different yeast species (Strucko et al., 2021). Moreover, the vector is equipped with a Cas9 gene expression cassette composed by the *K. phaffii TDH3p* (GAP1p) promoter, *cas9-NLS (SV40)* open reading frame adapted from (DiCarlo et al., 2013), and a *CYC1t* terminator from *S. cerevisiae*; and a USER cloning cassette (UCS) (Nour-Eldin et al., 2006) for effortless integration of desired sgRNA expression constructs (Figure 1**a**) encoding one or more guiding sequences.

**Figure 1.**
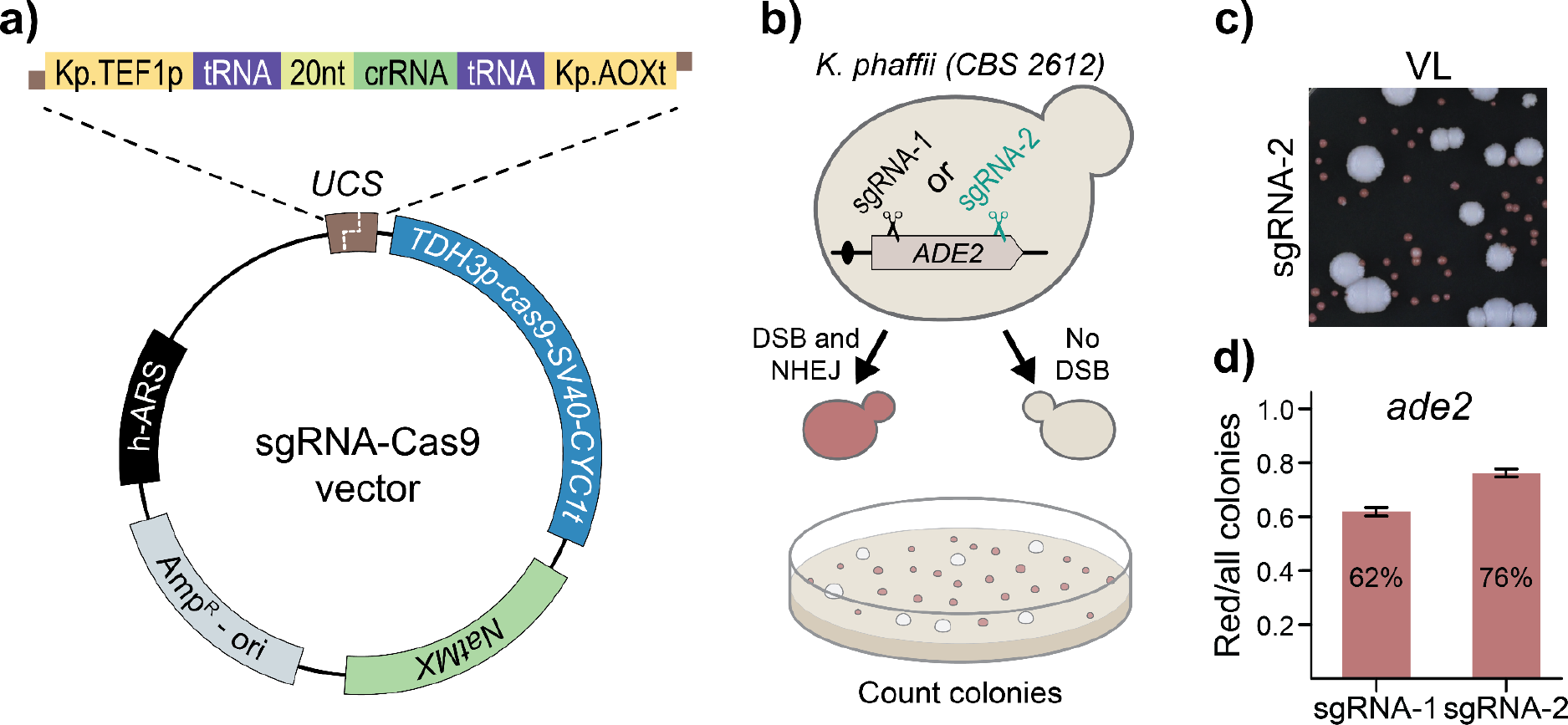
The design of the CRISPR-Cas9 system for *K. phaffii*. **a)** A schematic representation of the sgRNA-cas9 expression vector. UCS - USER cloning site. **b)** Screening setup to assess the functionality of our sgRNA-cas9 tool. Wild-type transformants from large white colonies while *ade2* transformants form small red/pink colonies. **c)** A snippet of a typical transformation plate in an NHEJ proficient *K. phaffii* strain transformed with the pDIV273 plasmid. VL - visible light. **d)** Ratios of red/total colonies obtained from two different experiments employing plasmids coding for two different sgRNA species, which target the *ADE2* gene in different positions as indicated in panel **b**. Error bars represent standard deviation based on four biological replicates.

Currently, several methods for expression of sgRNA in *K. phaffii* are available, but the most common ones rely on RNA Pol-II based expression cassettes equipped with self-cleaving ribozyme sequences for liberation of the functional sgRNA (Liao et al., 2021; Weninger et al., 2016) or Pol-III cassettes flanked with tRNA sequences (Dalvie et al., 2020).

However, assemblies of the reported sgRNA expression cassettes are rather complicated; especially, when multiple (more than two) sgRNAs need to be expressed. To allow for simple sgRNA expression cassette construction, we adopted a system we originally developed for *Debaryomyces hansenii*, (Strucko et al., 2021). Specifically, the cassette consist of the Pol-II *TEF1p* promoter from *K. phaffii*, an sgRNA coding sequence flanked by heterologous tRNA-Gly sequences from *Aspergillus nidulans*, and a *AOXt* terminator from *K. phaffii* (see Figure 1**a** and Materials and Methods). By using USER fusion (Nour-Eldin et al., 2006), simple and complex gRNA expression cassettes can easily be introduced into the CRISPR vector as described in (Supplementary Figure S5) and (Nødvig et al., 2018; Strucko et al., 2021).

To assess the functionality of our design, we constructed two sgRNA-Cas9 expressing CRISPR plasmids *sgRNA1-ADE2* (pDIV273) and *sgRNA2-ADE2* (pDIV274) targeting two different sequences in the *ADE2* gene of *K. phaffii* (Figure 1**b** and Materials and Methods). In this experiment, we exploited that mutation of *ADE2* can be easily assessed by visual screening as *ade2* cells accumulate red pigment (Figure 1**c**), (Roman, 1956). Moreover, we chose this gene as a target to demonstrate robustness of our method. Hence, since *ADE2* disruption in *K. phaffii* is accompanied by a significant fitness loss (Du et al., 2012); transformants that escape CRISPR mutagenesis will prevail in case CRISPR mutagenesis is inefficient. The two plasmids were independently transformed into *K. phaffii* to test whether they were able to induce sgRNA-Cas9-mediated mutations via erroneous NHEJ DNA repair. In both cases, red and white transformants were formed (Supplementary Figure S6). The fractions of red transformants (Figure 1**d**) in the two experiments were similar (62% to 76%, respectively), and were comparable with previously reported efficiencies (Liao et al., 2021). Lastly, Sanger sequencing of three random red colonies from each experiment that all *ade2* mutants investigated *ADE2* gene contained disruptive indels sites (the likely result of erroneous NHEJ DNA repair) at the Cas9 cleavage sites (See Supplementary Figure S7). We conclude that our CRISPR method can be used to efficiently generate gene mutations even in a case where the mutant products grow poorly compared to the wild-type strain.

### CRISPR-Cas9 induced gene targeting using linear DNA fragments

*K. phaffii*, like many other non-conventional yeasts and higher eukaryotes, prefers to insert DNA received by transformation into the genome via the NHEJ pathway. To increase the gene-targeting efficiency, it may therefore be an advantage to eliminate the NHEJ pathway; and for *K. phaffii*, it has been demonstrated that gene targeting efficiencies are dramatically improved in strains containing a disrupted *KU70* homolog (Näätsaari et al., 2012; Wang et al., 2023). In addition, NHEJ inactivation enables efficient use of the TAPE tool to assess whether specific sgRNA species are able to facilitate Cas9 cleavage at the target sites (Nødvig et al., 2018; Vanegas et al., 2017). For these reasons, we constructed an NHEJ-deficient strain (sDIV360) by inactivating *KU70* via single nucleotide deletion (A32Δ) resulting in a frameshift mutation and a stop codon (see Materials and Methods and Supplementary Figure S4). A subsequent TAPE experiment analyzing *sgRNA1-ADE2* and *sgRNA2-ADE2* plasmids confirmed that the NHEJ pathway was inactivated and both sgRNAs were efficiently guiding Cas9 nuclease to the intended locus (see Supplementary Figure S8).

Next, we examined whether our CRISPR-based method allowed for efficient gene targeting. As a proof of concept, we have decided to integrate two gene-targeting substrates (GTSs), both expressing yellow fluorescent protein (mVenus), into two sites located on two different chromosomes of *K. phaffii*. The integration sites (IS) were positioned in the intergenic genetic regions (Figure 2**a**) between transcriptionally active genes (Liang et al., 2012) to ensure high expression and at the same time minimize accompanying fitness defects. In addition, we tested GTSs containing long ∼1kb (LH) and short 60bp (SH) homology arms to target the GTS to each of the two integration sites. Moreover, to address the effect of NHEJ, we co-transformed sgRNA-Cas9 plasmids and the corresponding GTSs into NHEJ-proficient (WT) and -deficient (*ku70*) strain backgrounds (Figure 2). When we examined the fluorescence of the transformants obtained with wild-type strains after transformation with the two different GTSs into the two different integration sites, we found that most were fluorescent. For targeting of the GTS containing long homology arms into IS1 and IS3, we found that 90% or 74% of the transformants were fluorescent, respectively. A similar result was obtained in the targeting experiment employing GTS containing long homology arms. Diagnostic PCRs of sets of randomly selected transformants, from the four different experiments, testing for the presence of mVenus in the integration sites produced numbers that closely matched the numbers of the fluorescent colonies obtained from the corresponding experiments. Hence, although GTSs integrate efficiently into both integration sites, the efficiency was slightly higher for the setup targeting IS1. Importantly, the same numbers were obtained by using either long or short homology arms at both IS1 and IS3 showing that no advantage was gained by extending the length of the homology arm from 60 to 1000 bps. Further inspection of the fluorescence levels of the transformants revealed large clonal differences indicating that wild-type stains likely contain multiple copies of the GTSs (see Figure 2**b** and Supplementary Figure S9). Additional copies may be the result of NHEJ mediated polymerization of GTSs prior to incorporation into the integration site and/or NHEJ mediated integration of GTSs into other random sites in the genome. In agreement with this view, *ku70* strains produced transformants that all produced yellow fluorescence at the same level (Figure 2**c** and Supplementary Figure S9). Moreover, the level of the fluorescence obtained with *ku70* strains matched the lower levels obtained for transformants obtained with wild-type strains indicating that a single mVenuscopy has been inserted into the desired integration site by HR in the *ku70* strains. The gene-targeting efficiency exceeded 95% for all four experiments, and like with wild-type cells, no advantage was achieved by adding the longer homology arms. Finally, our experiments suggest that CRISPR mediated gene targeting may occur more robustly in the absence of NHEJ since the setup for targeting *mVenus* into IS3 works suboptimally in wild-type cells but is highly efficient in *ku70* strains (Figure 2**d** and **e** and Supplementary Figure S10 and S11).

**Figure 2.**
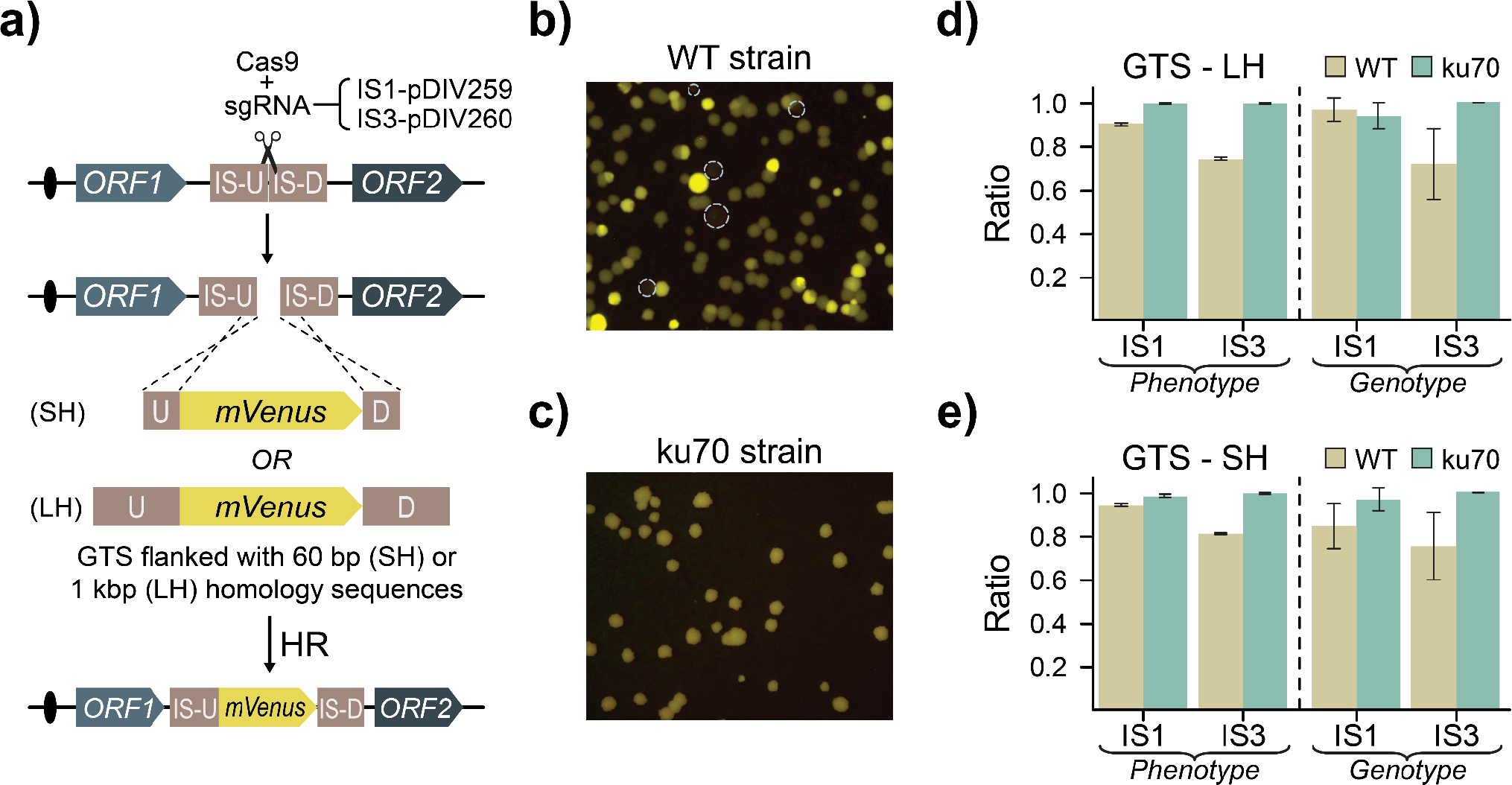
Gene integration strategy using CRISPR-Cas9 tool. **a)** Schematic depiction of gene targeting using intergenic chromosomal regions. GTS - gene targeting substrate, IS - integration site, SH - short homology, LH - long homology sequence, U - upstream and D - downstream. **b)** A qualitative snippet of a typical transformation plate detecting for yellow fluorescence light emitted by NHEJ proficient strains. Dashed circles show the location of non-fluorescent colonies. **c)** A qualitative snippet of a typical transformation plate detecting NHEJ deficient strains emitting yellow fluorescence light. All colonies appear to have similar fluorescence intensities. **d)** Results of gene targeting efficiencies using 1kb long targeting sequences and **e)** using short targeting sequences in integration sites IS1 and IS3. Phenotype - results show the ratio of fluorescent colonies vs. total colonies observed on transformation plates (Note that in WT- background strains colonies of different fluorescence intensities were included in calculation). Genotype - results show the ratio of colonies with correctly integrated GTS vs. total number of tested colonies for the corresponding GTS. Error bars represent standard deviation based on four biological replicates.

### Single-stranded DNA oligonucleotides are efficient gene targeting substrates

In metabolic engineering experiments, it is often necessary to rewire the host’s native metabolism by multiple gene deletions and/or specific point mutations. In *K. phaffii*, this is typically achieved by using linear double stranded GTSs constructed by PCR (Dalvie et al., 2022), in steps, which are time consuming and costly in large scale experiments. In the yeast *S. cerevisiae*, short double-stranded oligonucleotides are successfully applied for gene editing in CRISPR experiments (DiCarlo et al., 2013). Conversely, single-stranded oligonucleotides are used for gene editing in filamentous fungi (Nødvig et al., 2018; Vanegas et al., 2019) and the non-conventional yeast *D. hansenii* (Strucko et al., 2021). Thus, we investigated whether short single-stranded DNA can be applied as GTS in *K. phaffii*. For this purpose, we designed single-stranded 90-mers (ss-GTS) that are able to either introduce a stop codon (PR1292) or completely delete the *ADE2* gene (PR1291) (Figure 3**a** and **b**). We performed experiments in both WT and *ku70* strain backgrounds by co-transforming specific CRISPR plasmids encoding Cas9-sgRNA nucleases that are able to cleave *ADE2* at strategic places (see Figure 3) with the relevant ss-GTSs as repair template. In all cases, *ADE2* was mutated very efficiently as the ratio of red to total number of transformants varied from 92% to 95% in (Figure 3**c** and **d**). We next genotyped randomly selected red colonies (N=32) from each experiment to assess whether single-stranded oligonucleotide-guided genetic modifications were introduced as intended (see Materials and Methods and Supplementary Figure S12). When the mutagenic stop-codon oligonucleotide (PR1292) was used, 97% of analyzed red colonies contained correct edits in WT background as compared to 100% in *ku70* background (Figure 3**c**). When the gene-deletion oligonucleotide (PR1291) was applied, 31% and 100% of the red colonies contained the expected gene deletion in WT and *ku70* strains, respectively (Figure 3**d**).These results demonstrate that short single-stranded oligonucleotides can be efficiently used as GTSs for precise CRISPR based gene editing in *K. phaffii*, in particular if the NHEJ pathway is inactivated.

**Figure 3.**
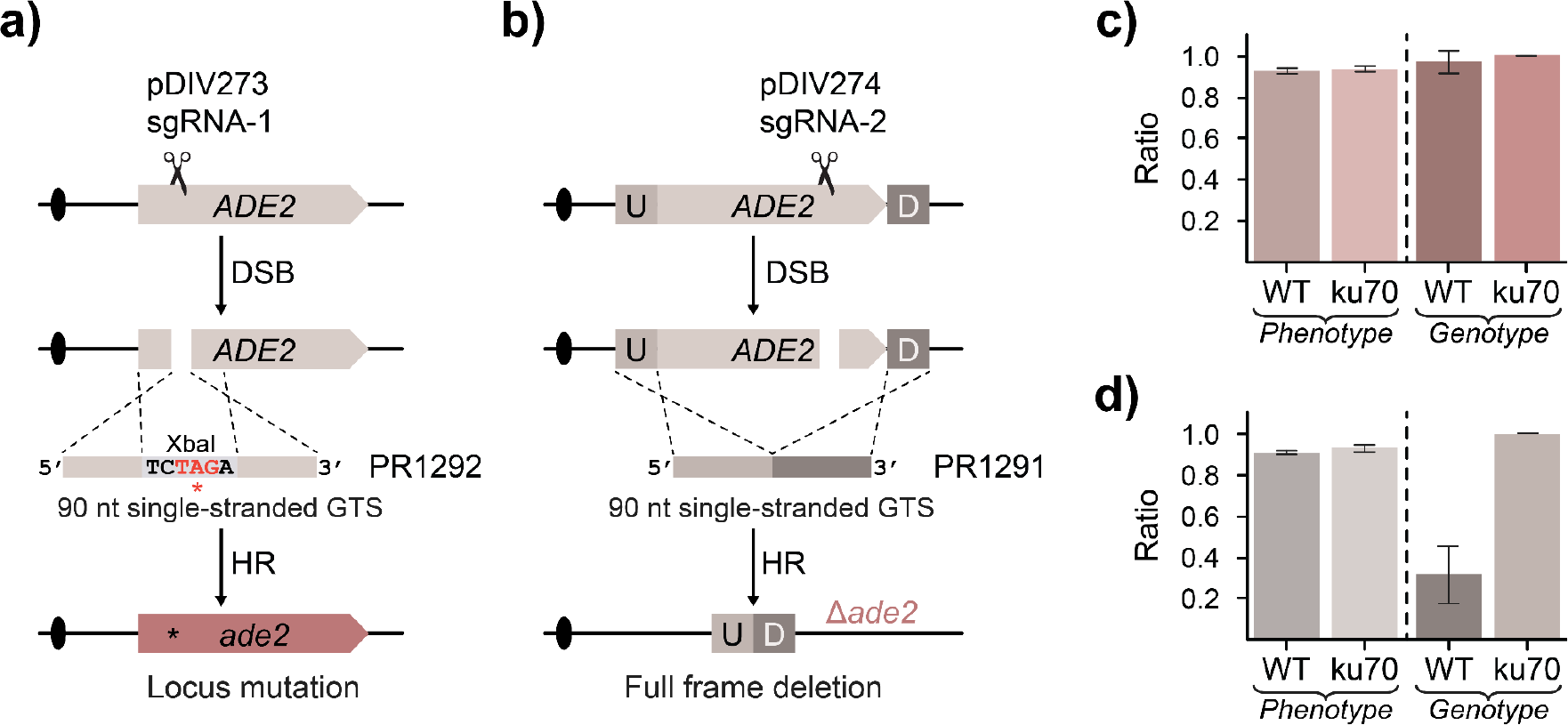
Single-stranded oligonucleotide based gene editing in *K. phaffii*. **a)** Schematic depiction of site specific mutagenesis using short single-stranded gene targeting substrate (GTS). DSB - double strand break, HR - homologous recombination. The oligonucleotide PR1292 contains an XbaI recognition site designed to introduce an in-frame stop codon. **b)** Schematic depiction of site specific mutagenesis using short single-stranded GTS (PR1291) for full gene deletion. Error bars represent standard deviation based on four biological replicates. U - upstream and D - downstream. **c)** Results of *ADE2* locus mutation efficiencies using short oligonucleotide. The bars show the ratio of red colonies with correctly integrated GTS vs. total number of tested red colonies for the corresponding GTS in *ku70* - NHEJ deficient and *WT* - NHEJ proficient strain background. Error bars represent standard deviation based on four biological replicates. **d)** Results of *ADE2* deletion efficiencies using short oligonucleotide. The bars show the ratio of red colonies with correctly integrated GTS vs. total number of tested red colonies for the corresponding GTS in *ku70* - NHEJ deficient and *WT* - NHEJ proficient strain background. Error bars represent standard deviation based on four biological replicates.

### Visual marker system for easy distinction of NHEJ-deficient/proficient strains

Disruption of NHEJ pathway in *K. phaffii* and higher eukaryotes improves gene targeting and reduces unwanted genome-wide mutation in CRISPR-based experiments (Garrigues et al., 2023; Schusterbauer et al., 2022). However, the presence of the *ku70* mutation may influence the final phenotype of the engineered strain. For example, deletion of *YKU70* in *S. cerevisiae* confers a temperature-sensitive phenotype, affects telomere length, and expression of telomere-proximal genes (Fellerhoff et al., 2000). Moreover, it was reported that the *ku70* genotype confers an increased UV-sensitivity phenotype in *K. phaffii* (Näätsaari et al., 2012). It is therefore advisable to restore the NHEJ pathway in CRISPR engineered strains prior to functional characterization and data interpretation. To facilitate this routine, we have developed a transient *KU70* gene disruption system that allows visual distinction between NHEJ-deficient and -proficient cells. Specifically, NHEJ was eliminated by inserting the common color-marker gene *uidA* gene (Jefferson et al., 1987) into the ORF of *KU70* to disrupt its gene function. As a result, *ku70* strains can therefore easily be identified on media supplemented with X-Gluc as they turn blue, Materials and Methods. Once desirable modification(s) have been introduced into the *ku70::uidA* strain, the NHEJ pathway can be easily restored in a CRISPR event by using a Cas9/sgRNA nuclease targeting the *uidA* cassette and a short PCR derived DNA repair fragment that allows the *KU70* gene to be restored. Subsequent plating on X-Gluc media readily identifies *KU70* strains as they turn white (Figure 4**a**).

**Figure 4.**
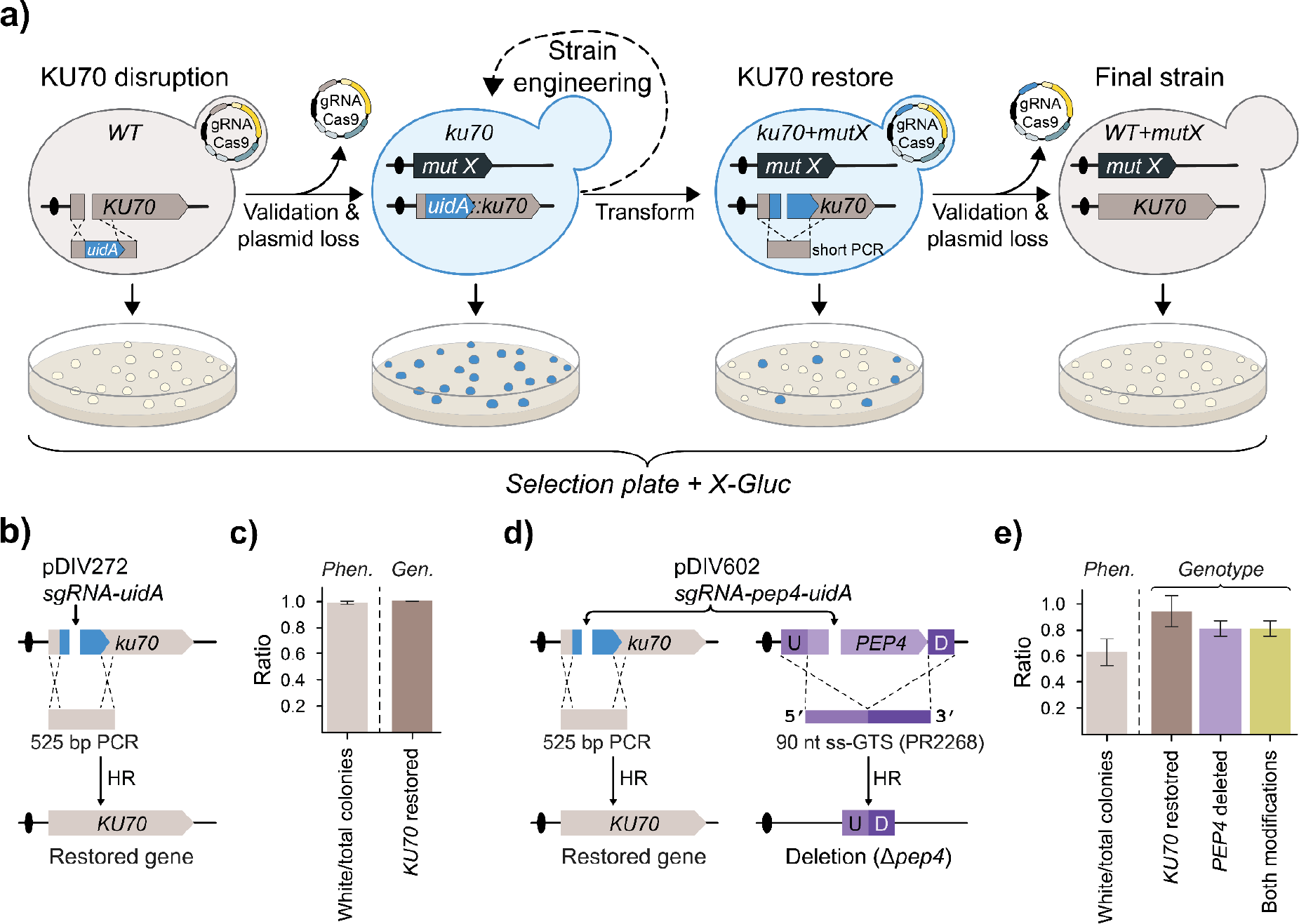
Color-based system for deactivation and reactivation of *KU70* gene in *K. phaffii*. **a)** A schematic illustration depicting the steps of *KU70* gene disruption by *uidA* cassette, its genetic manipulation and *KU70* reversion to wild-type. Petri dishes illustrate expected colony phenotypes on media supplemented with X-Gluc and mutX represents a desired mutation(s) performed during genetic engineering of the yeast. **b)** Experimental setup to restore disrupted *KU70* gene using short PCR-based GTS. **c)** Results of *KU70* gene restoration. Light brown - the ratio of white colonies vs. total colonies observed on transformation plates, dark brown - the ratio of colonies with correctly restored *KU70* vs. total number of tested white colonies. Error bars represent standard deviation based on four biological replicates. **d)** Multiplex experimental setup for simultaneous deletion of *PEP4* gene and *KU70* restoration. U - upstream and D - downstream. **e)** Results of an experiment from panel (d). Light brown - the ratio of white colonies vs. total colonies observed on transformation plates, dark brown - the ratio of colonies with correctly restored *KU70* vs. total number of tested white colonies, purple - the ratio of colonies with correctly integrated GTS at *PEP4* locus vs. total number of tested white colonies, and yellow - the ratio of colonies that have both modifications validated vs. total number of tested white colonies for both modifications. Error bars represent standard deviation based on three biological replicates.

We tested the functionality of our system by co-transforming the *ku70::uidA* (sDIV291) strain with sgRNA-uidA (pDIV272) plasmid and a PCR fragment harboring the restoring sequence of *KU70* gene (Figure 4**b**). In this experiment, virtually all (∼98%) of transformed cells formed white colonies on X-Gluc media (Figure 4**c** and Supplementary Figure S13). The genotyping of 32 randomly selected white colonies revealed that in 100% of cases the *KU70* gene was restored to the wild-type sequence (Figure 4**c** and Supplementary Figure S14). Based on the latter results, we conclude that our system offers quick and efficient reversion of *ku70* to wild-type. Next, we asked whether our *ku70::uidA* system can be applied in multiplex experiments involving gene editing of a desired locus and simultaneous reversion of the edited strain to NHEJ proficient state. As a proof-of-concept, we decided to delete the *PEP4* gene (coding for vacuolar aspartyl protease) and restore *ku70::uidA* locus to wild-type *KU70* in one round of transformation (see Figure 4**d**). We performed this experiment by co-transforming the NHEJ-deficient (sDIV291) strain with a plasmid pDIV602 (expressing two sgRNAs targeting Cas9 to *PEP4* and *uidA*) and two repair templates, a PCR fragment to restore *KU70* and an ss-GTS (PR2268) to delete the *PEP4* gene. Transformants were successfully obtained, but the number of colonies per transformation were noticeably lower as compared to the previous experiments where only *ku70* locus was targeted (Supplementary Figure S13). Moreover, only ∼40% of the transformants displayed the blue phenotype on X-Gluc plates (Figure 4**e** and Supplementary Figure S13). Based on this result, we note that the efficiency of a specific gene-editing event appears reduced when it is part of a multiplexing experiment as compared to the efficiency obtained in an experiment where it is introduced as the only modification. Next, to examine for the overall multiplexing efficiency, we genotyped 20 randomly selected white colonies on X-Gluc media (see Materials and Methods, and Supplementary Figure S14). This analysis showed that ∼92% of the transformants contained a restored *KU70* gene and 80 % contained the *pep4* genotype (Figure 4**e**). However, when we examined the transformants that were NHEJ they were all *pep4* showing that cells that are proficient for CRISPR gene editing are capable of introducing two modifications at high efficiency. The latter suggests that the colony color itself can therefore serve as an indicator for selection of cells that most likely underwent the intended genetic modification(s), in this case a *PEP4* deletion and *ku70* reversion.

## Concluding remarks

In this study, we have developed a CRISPR-Cas9 based method that can serve as an improved addition to the currently available gene engineering tool-box for *K. phaffii*. We have demonstrated that our CRISPR tools can be applied for editing in either NHEJ-proficient or -deficient strain backgrounds. Additionally, our results strongly indicate and support multiple previous studies showing that NHEJ-deficient strain backgrounds are beneficial for gene-editing experiments. Importantly, we have demonstrated that short single-stranded oligonucleotides facilitate DNA DSB repair induced by CRISPR nuclease, and thus, can be used to introduce specific gene deletions or mutations with very high efficiency. Successful application of short single-stranded oligonucleotides will dramatically reduce the cost and time associated with construction of GTSs in large scale experiments. Lastly, we address the widely ignored fact that non-functional *KU70* gene is associated with unwanted trade-offs that can potentially compromise the performance of *K. phaffii* in physiological characterizations as well as a cell factory. Thus, we devised a color based method that allows for transient color marker based inactivation of *KU70* and successfully demonstrated its application in genetic engineering. Lastly, we envision that our color marker based system could serve as a highly useful add-on that can be easily implemented into automated strain construction workflows.

## Author contributions

## Acknowledgements

We thank Irina Borodina for kindly providing the pCfB2312 plasmid.

