## Supplementary File for "Oligonucleotide-based CRISPR-Cas9 toolbox for efficient engineering of *Komagataella phaffii*"

### List of Supplementary Tables

Supplementary Table S1. Primers used in this study

Supplementary Table S2. Plasmids used in this study

### List of Supplementary Figures

Supplementary Figure S1. USER cloning workflow for assembly of Cas9-sgRNA vectors used in this study.

Supplementary Figure S2. USER cloning workflow for assembly of vectors harboring GTSs used in this study.

Supplementary Figure S3. PCR-based for assembly of linear GTSs used in this study.

Supplementary Figure S4. An excerpt of Sanger sequencing results of KU70 locus

Supplementary Figure S5. A generalized protocol for Cas9-sgRNA vector assembly using USER cloning.

Supplementary Figure S6. Images of Transformation plates of ADE2 experiment.

Supplementary Figure S7. An excerpt of  Sanger sequencing results of ADE2 locus.

Supplementary Figure S8. Images of agar plates representing a TAPE experiment.

Supplementary Figure S9. Images of agar plates with yellow fluorescent colonies.

Supplementary Figure S10. Validation of integrated mVenus cassette into integration site (IS1).

Supplementary Figure S11. Validation of integrated mVenus cassette into integration site (IS3).

Supplementary Figure S12. Validation of modified ADE2 locus using short single stranded DNA oligonucleotides as GTS.

Supplementary Figure S13. Images of agar plates of ku70 reversion and PEP4 deletion experiment.

Supplementary Figure S14. Validation of modified KU70 locus PEP4 gene deletion.

**Supplementary Table S1**.Primers used in this study.

| Name | Sequence | Purpose |
|---|---|---|
| PR_DIV0008 | ATCTGTCAUATGTTACGTCCTGTAGAAACCCC | For construction of pDIV033 plasmid |
| PR_DIV0010 | CGTGCGAUGCACACACCATAGCTTCAAAATG | For construction of pDIV033 plasmid |
| PR_DIV0011 | ATGACAGAUTTTGTAATTAAAACTTAGATTAGATTGCT | For construction of pDIV033 plasmid |
| PR_DIV0105 | CACGCGAUCTTCGAGCGTCCCAAAACC | For construction of pDIV033 plasmid |
| PR_DIV0120 | ATCTGTCAUATGGTTAGTAAGGGGGAAGAGCT | For construction of pDIV479 plasmid |
| PR_DIV0681 | CGTGCGAUCGCGTGCATTCTTTTTGTAGAAATGTCTTGGTGT | For construction of the basic Cas9-expressing vector |
| PR_DIV0682 | ATTGTGUTTTGATAGTTGTTCAATTGATTGAA | For construction of the basic Cas9-expressing vector |
| PR_DIV0683 | ACACAAUGGACAAGAAGTACTCCATTGG | For construction of the basic Cas9-expressing vector |
| PR_DIV0684 | CACGCGAUGTACCGGCCGCAAATTAA | For construction of the basic Cas9-expressing vector |
| PR_DIV0810 | CGCCAGATGTAATCACCGC | Primers for validation of KU70 Locus |
| PR_DIV0811 | CATGGCGAAGACTCATAAGC | Primers for validation of KU70 Locus |
| PR_DIV0812 | CTTCCCTGGCTTCCATGG | Primers for validation of KU70 Locus |
| PR_DIV0816 | GGCATCCTACTGCTAATCCTTC | For validation of PEP4 locus |
| PR_DIV0817 | GGCAATTGACATCGTAGTACC | For validation of PEP4 locus |
| PR_DIV0818 | GCTCCTTAACATCCTCTCCATG | For validation of PEP4 locus |
| PR_DIV0819 | CGTGCGAUTAACTCTGGCAACCAGTAACACG | For construction of the basic tRNA-based sgRNA expression cassette |
| PR_DIV0820 | ATCCGTGAAUCGAACACGGGCCTCATCGATGGCAACGATGAATTCTACCACTAGACCAATGATGCGTTGGCGAATAACTAAAATGTATGT | For construction of the basic tRNA-based sgRNA expression cassette |
| PR_DIV0821 | ATTCACGGAUGATGCAACGACGACTGTTTGCGCTTAGTTTTAGAGCTAGAAATAGCAAGTTAAAATAAG | For construction of the basic tRNA-based sgRNA expression cassette |
| PR_DIV0822 | ATCCTCTTGAUGCATCATCCGTGAATCGAAC | For construction of the basic tRNA-based sgRNA expression cassette |
| PR_DIV0823 | aTCAAGAGGAUGTCAGAATGCCATTTGCC | For construction of the basic tRNA-based sgRNA expression cassette |
| PR_DIV0824 | CACGCGAUTCTGTACTCTGAAGAGGAGTGGG | For construction of the basic tRNA-based sgRNA expression cassette |
| PR_DIV0961 | GGTCTTAAUTAATGCCTCAGCTTCGAGCGTCCCAAAACC | For construction of pDIV479 plasmid |
| PR_DIV0966 | ATTCACGGAUGATGCAGAAACAGTCCAACAGTAAGCGTTTTAGAGCTAGAAATAGCAAGTTAAAATAAG | For construction of the tRNA-based sgRNA expression cassette (IS1) |
| PR_DIV0967 | GGGTTTAAUGGCTTGTCTGCCAATACTACT | For construction of GTSs with long homology (LH) sequences targeting integration site IS1 |
| PR_DIV0968 | GGACTTAAUAGTCGATTTTACGACTACCACA | For construction of GTSs with long homology (LH) sequences targeting integration site IS1 |
| PR_DIV0969 | GGCCGAAACCTCTCGTG | Validation primer for integration site IS1 |
| PR_DIV0970 | GGCATTAAUGGAAGCACATGGCCCTAC | For construction of GTSs with long homology (LH) sequences targeting integration site IS1 |
| PR_DIV0971 | GGTCTTAAUCCTGAATCAGCCTATGGGTC | For construction of GTSs with long homology (LH) sequences targeting integration site IS1 |
| PR_DIV0972 | CCTCCTGCTTCACTACCTC | Validation primer for integration site IS1 |
| PR_DIV0973 | ATTCACGGAUGATGCATGTTGATAATGGACCGTGGGGTTTTAGAGCTAGAAATAGCAAGTTAAAATAAG | For construction of the tRNA-based sgRNA expression cassette (IS3) |
| PR_DIV0974 | GGGTTTAAUGCAGTGGTTGGAGGAAGAG | For construction of GTSs with long homology (LH) sequences targeting integration site IS3 |
| PR_DIV0975 | GGACTTAAUCGGAGGATTTTGATAGAAATTGGC | For construction of GTSs with long homology (LH) sequences targeting |

| Name | Sequence | Purpose |
|---|---|---|
| | | integration site IS3 |
| PR_DIV0976 | CAGTTGCAGCAACAACCTC | Validation primer for integration site IS3 |
| PR_DIV0977 | GGCATTAAUTCAACACCAGCAGCATTCA | For construction of GTSs with long homology (LH) sequences targeting integration site IS3 |
| PR_DIV0978 | GGTCTTAAUGTGCCTTTGGAGGCTGG | For construction of GTSs with long homology (LH) sequences targeting integration site IS3 |
| PR_DIV0979 | CGTGAGAGGCTGATAGCC | Validation primer for integration site IS3 |
| PR_DIV1045 | ATGACAGAUGTTGGCGAATAACTAAAATGTATGTAG | For construction of pDIV479 plasmid |
| PR_DIV1253 | CCATATTAACATAACATGTATATAAACGTC | Standard primers for plasmid assembly validations and SANGER sequencing |
| PR_DIV1254 | GAAACCATTATTATCATGACATTAACC | Standard primers for plasmid assembly validations and SANGER sequencing |
| PR_DIV1255 | CGAAGTTATATTAAGGGTTGTCGAC | Standard primers for plasmid assembly validations and SANGER sequencing |
| PR_DIV1277 | GGGTTTAAUTTCACCTACGGGTCTGACTACC | Primers for Construction of pDIV643 plasmid |
| PR_DIV1279 | AAAGCATUGCGCACACACCATAGCTTCAAAATG | Primers for Construction of pDIV643 plasmid |
| PR_DIV1280 | ACACCTUCGAGCGTCCCAAAACCTTC | Primers for Construction of pDIV643 plasmid |
| PR_DIV1283 | ATTCACGGAUGATGCAAAGCAATACGACATCCACGAGTTTTAG AGCTAGAAATAGCAAGTTAAAATAAG | For construction of the basic tRNA-based sgRNA expression cassette (Ku70) |
| PR_DIV1285 | ATTCACGGAUGATGCATACTTCAGAAGCGTTGGACCGTTTTAG AGCTAGAAATAGCAAGTTAAAATAAG | For construction of the basic tRNA-based sgRNA expression cassette (uidA v2) |
| PR_DIV1286 | CGACGAGCCAGCACTTTATAG | Validation primers for ADE2 locus |
| PR_DIV1287 | GCCTCTTCCATCATAAGCCATAG | Validation primers for ADE2 locus |
| PR_DIV1288 | GGAGGACAAATTTCTACCGAAC | Validation primers for ADE2 locus |
| PR_DIV1289 | ATTCACGGAUGATGCAGCTGCGCAAGACCACATCGAGTTTTA GAGCTAGAAATAGCAAGTTAAAATAAG | For construction of the basic tRNA-based sgRNA expression cassette (ADE2 v1) |
| PR_DIV1290 | ATTCACGGAUGATGCACAATGGAGACCGAAGTGTTGGTTTTA GAGCTAGAAATAGCAAGTTAAAATAAG | For construction of the basic tRNA-based sgRNA expression cassette (ADE2 v1) |
| PR_DIV1291 | CTTTTCCAAGAATCGTAGAAACGATTAAAAAACTTCCAAACTCT CGAATTTAGTATTGTTTTTTAATAGATGTATATATAATAGTACACG | ss-GTS for ADE2 deletion |
| PR_DIV1292 | TTTTTCACCTGCTAAGCACATTAATGCTGCGCAAGACCACATCt AgATCATTCAAAGATGAGGAGGCTATCGCCAAGTTAGCTGCCA AAT | ss-GTS for ADE2 mutation |
| PR_DIV1509 | CGTAACAACAACTCACAGTCGATTGATTTGTAGACCTTTGTGG TAGTCGTAAAATCGACTGGTATTTGACAGGTTGGGGAG | For construction of GTS (60bp tails) integrating mVenus into IS1 |
| PR_DIV1510 | GAATGTTGTAAGAGGAGATTGGTTTTTAGGTGAGGAGAATGTA GGGCCATGTGCTTCCATCTTCGAGCGTCCCAAAACC | For construction of GTS (60bp tails) integrating mVenus into IS1 |
| PR_DIV1511 | GATCCAATGGGACAGGCTCCCCAAATTTATCAGCAACACGCC AATTTCTATCAAAATCCTGGTATTTGACAGGTTGGGGAG | For construction of GTS (60bp tails) integrating mVenus into IS3 |
| PR_DIV1512 | ATCACGGTGAGTCTCATATGAATAGTGCCCATTAGTGCGGACA GGTGAATGCTGCTGGTGCTTCGAGCGTCCCAAAACC | For construction of GTS (60bp tails) integrating mVenus into IS3 |
| PR_DIV1517 | GGGTTTAAUTAAGTCCTCAGCGGTATTTGACAGGTTGGGGAG | For construction of pDIV479 plasmid |
| PR_DIV2268 | ACATTCTAGAGTTCCATTTCTCAATTACTGATAATCAATTTAAAG GCAAGAATAAAAGTTGCTCAGCTGAACTTATTTGGTTACTTATC A | ss-GTS for PEP4 deletion |
| PR_DIV2269 | CGGGCATAACTTTAGGGATG | For validation of of integration in IS sites |
| PR_DIV2394 | AAAGTCCAUGGGTTTTAGAGCTAGAAATAGCAAGTTAAAATAA G | Assembly of multiplex sgRNA targeting PEP4 and uidA |
| PR_DIV2395 | ATGGACTTUCCTGAACCATGCATCATCCGTGAATCGAAC | Assembly of multiplex sgRNA targeting PEP4 and uidA |
| PR_DIV2396 | AGAAGCGTUGGACCGTTTTAGAGCTAGAAATAGCAAGTTAAAA TAAG | Assembly of multiplex sgRNA targeting PEP4 and uidA |
| PR_DIV2397 | AACGCTTCUGAAGTATGCATCATCCGTGAATCGAAC | Assembly of multiplex sgRNA targeting PEP4 and uidA |
| PR_DIV2449 | CGTGGAAGCAGCAACGAGG | Validation primes for restoring KU70 gene |

| Name | Sequence | Purpose |
|---|---|---|
| PR_DIV2451 | TGGCGATGATATCAAAGTTGG | Validation primes for restoring KU70 gene |
| PR_DIV2470 | AATGCTTUGGATGTCGTATTGCTTGCTGAC | Primers for Construction of pDIV643 plasmid |
| PR_DIV2471 | AAGGTGUCGGCTTCAGAAGGGAAATCT | Primers for Construction of pDIV643 plasmid |
| PR_DIV2472 | GGTCTTAAUAGGACAGGTGAAGTAAACCCG | Primers for Construction of pDIV643 plasmid |
| PR_DIV2599 | CTCTCTTGTAGGGGTCTCTACTGG | Validation primes for restoring KU70 gene |
| PR_DIV2660 | CGTGGAAGCAGCAACGAGG | For construction of GTS to restore KU70 |
| PR_DIV2661 | TTTCCAAATTCAGGTCTTGTAACTC | For construction of GTS to restore KU70 |

**Supplementary table S2.** Plasmids used this study.

| Name | Parent | Purpose | Source |
|---|---|---|---|
| pAC125 | --- | Backbone vector with USER cloning cassette (PacI/Nt.bbvCI) AmpR marker | Hansen et al. 2011 |
| pCfB2312 | --- | Expresses Cas9 cassette. Template for PCR for cloning pDIV151. | Stovicek et al 2015 |
| pDIV019 | --- | Multi-species compatible vector with USER cassette (AsiSI/Nb.BsmI) and NatMX marker | Strucko et al. 2021 |
| pDIV033 | pDIV019 | A template for the uidA cassette amplification and positive control for X-Gluc screen. | This study |
| pDIV151 | pDIV019 | Cas9 expressing vector harboring USER cloning cassette (AsiSI/Nb.BsmI) | This study |
| pDIV153 | pDIV151 | Generic Cas9-sgRNA vector to be used as a template for new plasmid cloning | This study |
| pDIV259 | pDIV151 | Cas9-sgRNA vector targeting IS1 locus | This study |
| pDIV260 | pDIV151 | Cas9-sgRNA vector targeting IS3 locus | This study |
| pDIV270 | pDIV151 | Cas9-sgRNA vector targeting KU70 gene | This study |
| pDIV272 | pDIV151 | Cas9-sgRNA vector targeting uidA gene locus 2 | This study |
| pDIV273 | pDIV151 | Cas9-sgRNA vector targeting ADE2 gene at locus 1 | This study |
| pDIV274 | pDIV151 | Cas9-sgRNA vector targeting ADE2 gene at locus 2 | This study |
| pDIV479 | pAC125 | Basic plasmid expressing mVenus cassette flanked with USER cloning sites | This study |
| pDIV505 | pDIV479 | Integrative plasmid ~1000 bp overhangs into Kp IS1 | This study |
| pDIV506 | pDIV479 | Integrative plasmid ~800 bp overhangs into Kp IS3 | This study |
| pDIV581 | pDIV151 | Cas9-sgRNA vector targeting PEP4 gene | This study |
| pDIV602 | pDIV151 | Multiplex Cas9-sgRNA vector targeting PEP4 and uidA genes | This study |
| pDIV643 | pAC125 | Integrative plasmid ~ 500 bp overhangs for disruption of KU70 gene via uidA cassette. | This study |

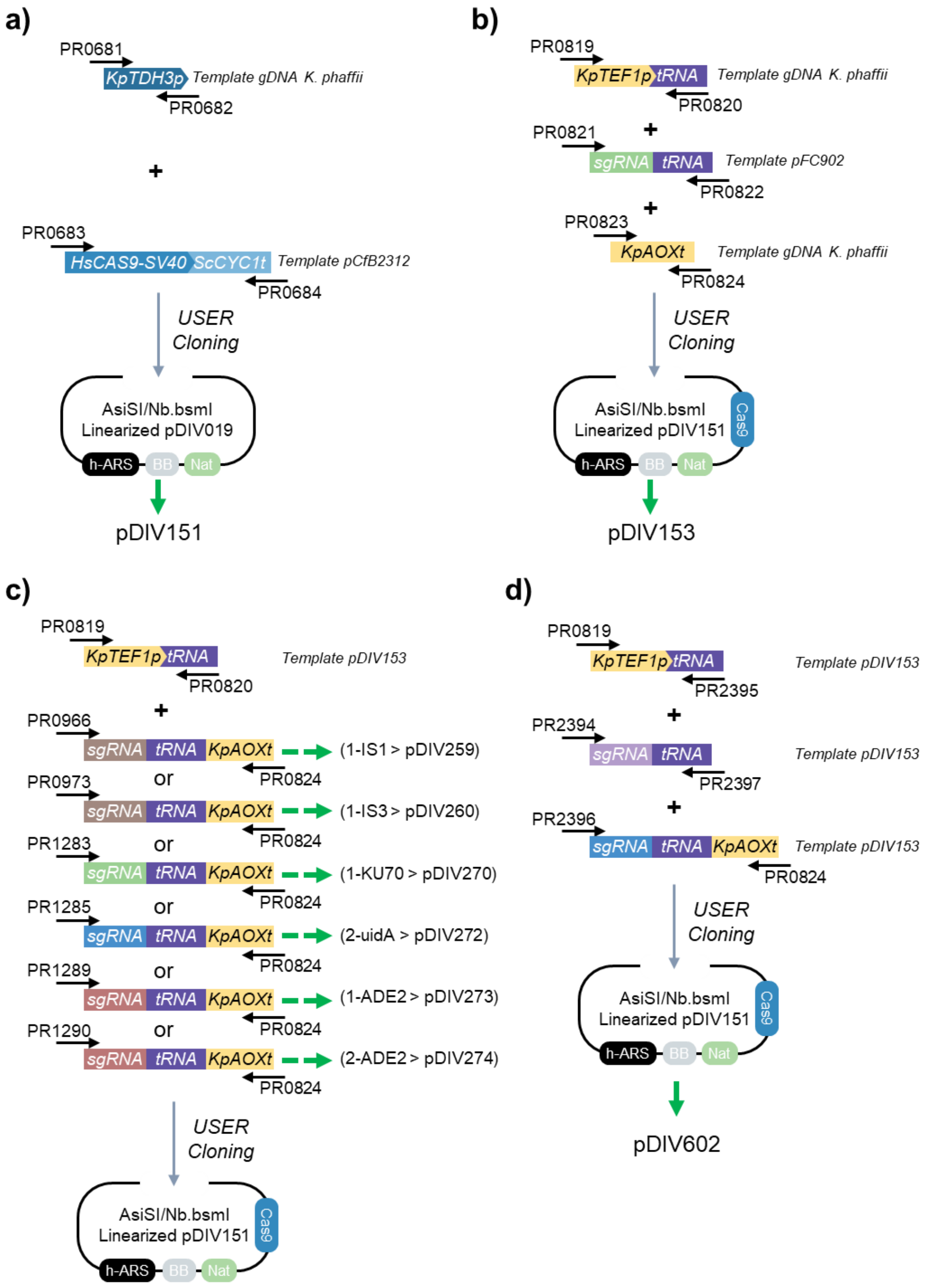


**Supplementary Figure S1.** USER cloning workflow for assembly of Cas9-sgRNA vectors used in this study. Cloning procedure for assembly of Cas9 expressing vectors **a)** pDIV151, **b)** pDIV153, **c)** pDIV259, pDIV260, pDIV270, pDIV272, pDIV273 and pDIV274, and **d)** pDIV602. Thin black arrows (annotated with PR####) represent primers used for PCR amplification. Colored text boxes show DNA fragments containing functional parts. Thick green arrows indicate the name of a final construct in each specific case.

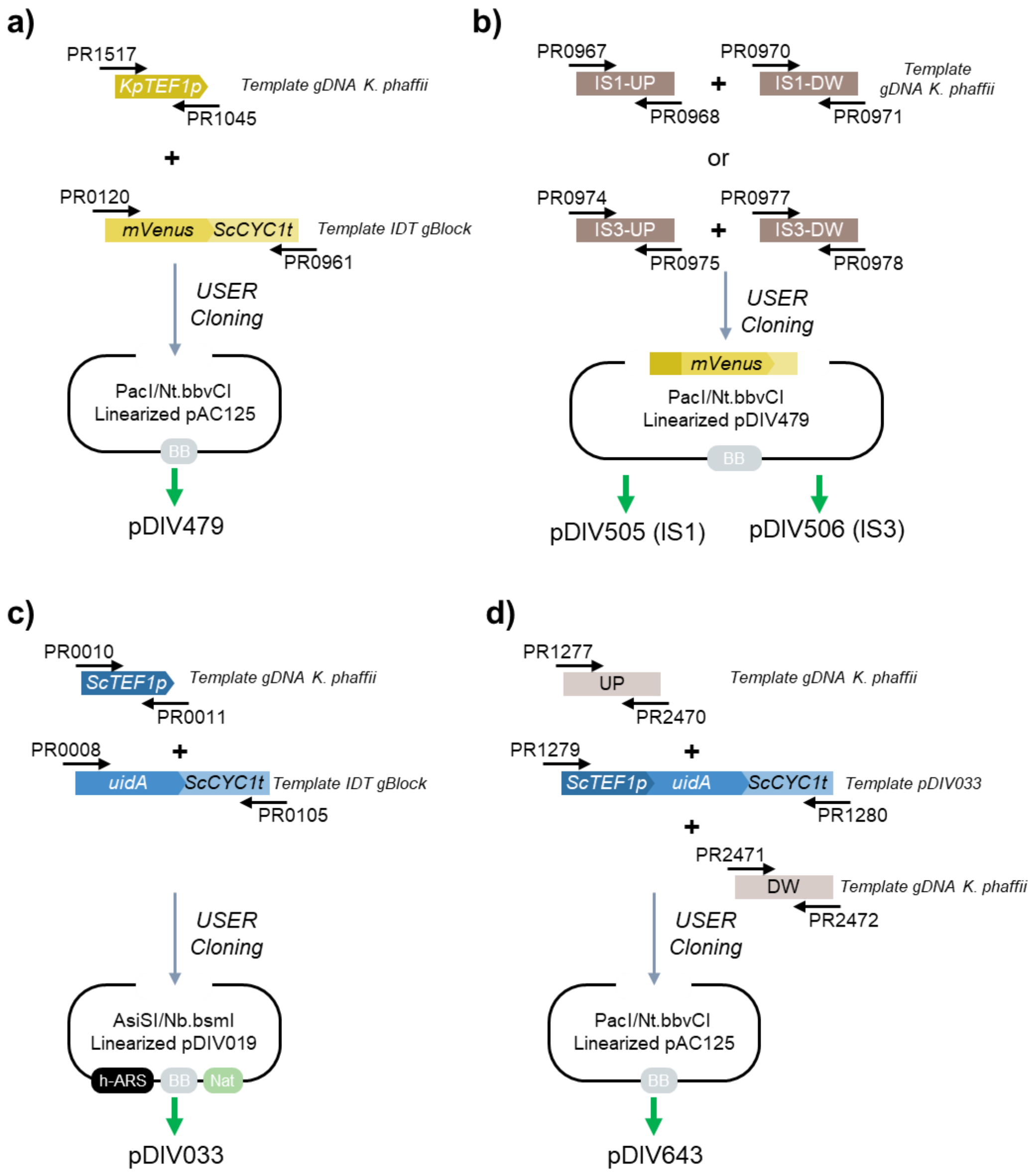


**Supplementary Figure S2.** USER cloning workflow for assembly of vectors harboring GTSs used in this study. Cloning procedure for assembly of **a)** pDIV479 plasmid, **b)** pDIV505 and pDIV506 plasmids, **c)** pDIV033, and **d)** pDIV643 plasmid. Thin black arrows (annotated with PR####) represent primers used for PCR amplification. Colored text boxes show DNA fragments containing functional parts. Thick green arrows indicate the name of a final construct in each specific case.

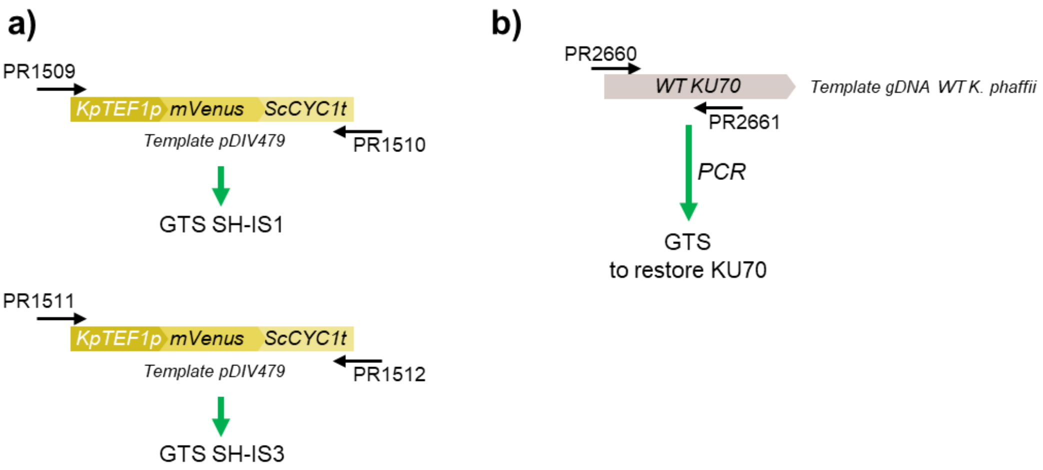


**Supplementary Figure S3.** PCR-based for assembly of linear GTSs used in this study. Amplification of PCR fragments **a)** GTS-SH-IS1 and GTS-SH-IS3 and **b)** GTS to restore KU70 gene. Thin black arrows (annotated with PR####) represent primers used for PCR amplification. Colored text boxes show DNA fragments containing functional parts. Thick green arrows indicate the name of a final construct in each specific case.

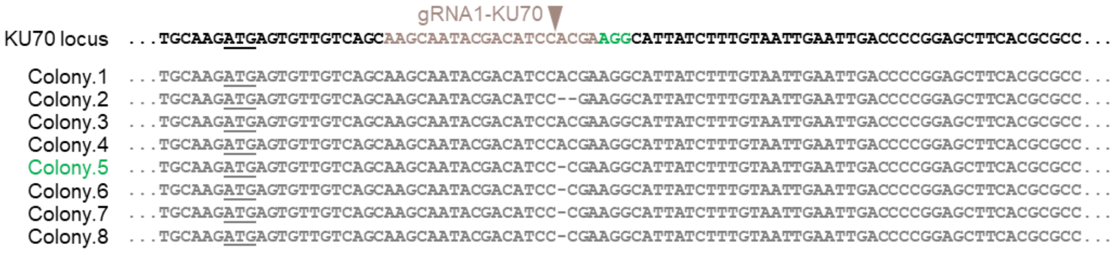


**Supplementary Figure S4.** An excerpt of Sanger sequencing results of *KU70* locus. Eight random colonies were selected from transformants by pDIV270 plasmid (sgRNA1-KU70). Light brown text indicates 20 nt guiding sequence, green denotes for PAM sequence. Underlined text indicated a start codon of the *KU70* gene.

a)

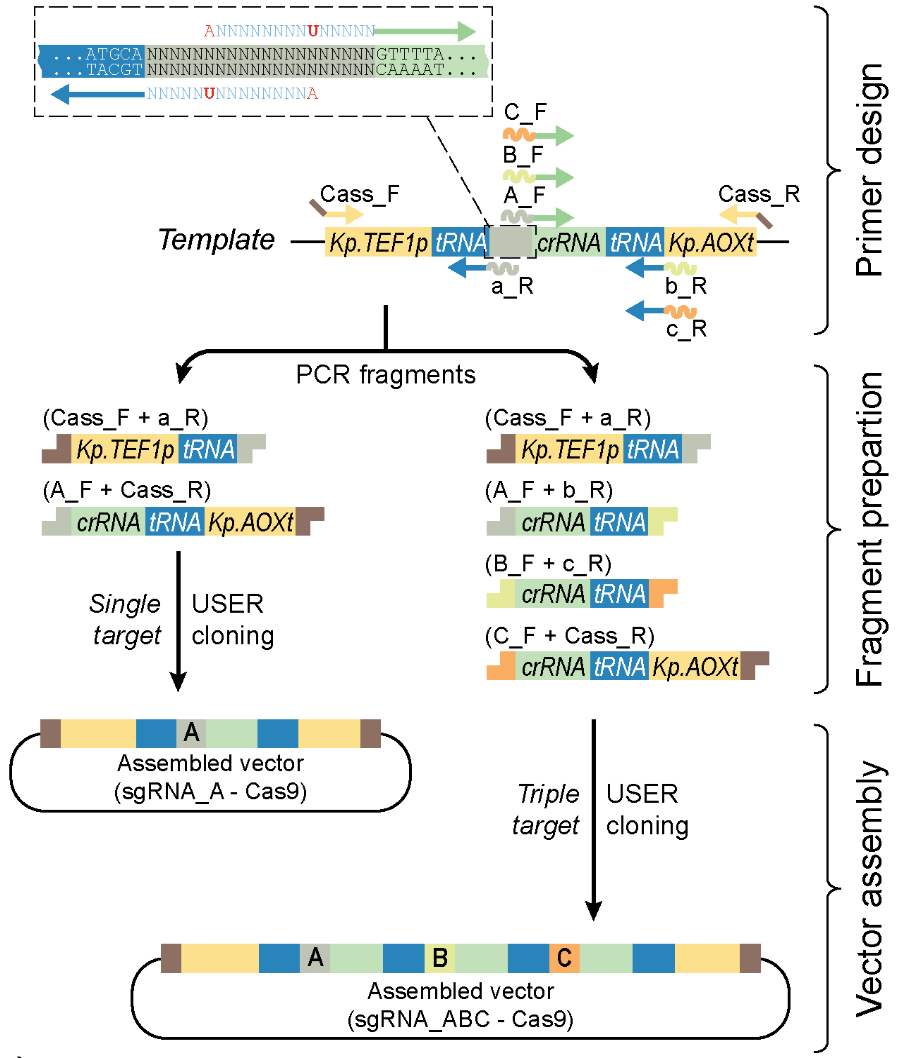


b)

| | |
|---|---|
| Cass_F | CGTGCGAUTAACTCTGGCAACCAGTAACACG |
| x_R | NNNNNNNNTGCATCATCCGTGAATCGAAC |
| X_F | NNNNNNNNGTTTTAGAGCTAGAAATAGCAAGTTAAAATAAG |
| Cass_R | CACGCGAUTCTGTACTCTGAAGAGGAGTGGG |

**Supplementary Figure S5.** A generalized protocol for Cas9-sgRNA vector assembly using USER cloning. **a)** Schematic depiction of a workflow for sgRNA-Cas9 vector assembly. The workflow can be split into three steps: 1-Primer design, 2-Fragment preparation, and 3-Vector assembly. In the step1, for each new sgRNA two primers (thick blue and green arrows) equipped with uracil containing tails are designed (depicted in the top left box). The tails are designed so that upon USER assembly it will encode a target specific 20nt guiding sequence (depicted as colorful wave lines and annotated with lowercase letters “a, b, c” for the reverse “R” primers, and capital letters “A, B, C” for forward “F” primers. In the step2, specific uracil tails containing fragments are generated by PCR using primers depicted in parenthesis. In case of a single sgRNA that would guide to the target “A”, two PCR fragments are generated and assembled into a Cas9 expressing vector (left side of the figure). In case of multiple sgRNAs that would target (in this example three targets) “A, B and C”, four fragments are generated and cloned into a Cas9 expressing vector (right side of the figure). Using the same logic, vectors expressing two, four or more sgRNAs can be assembled **b)** Standard primer sequences for sgRNA cassette cloning. N - represents a USER tail that would result in a unique targeting sequence upon assembly.

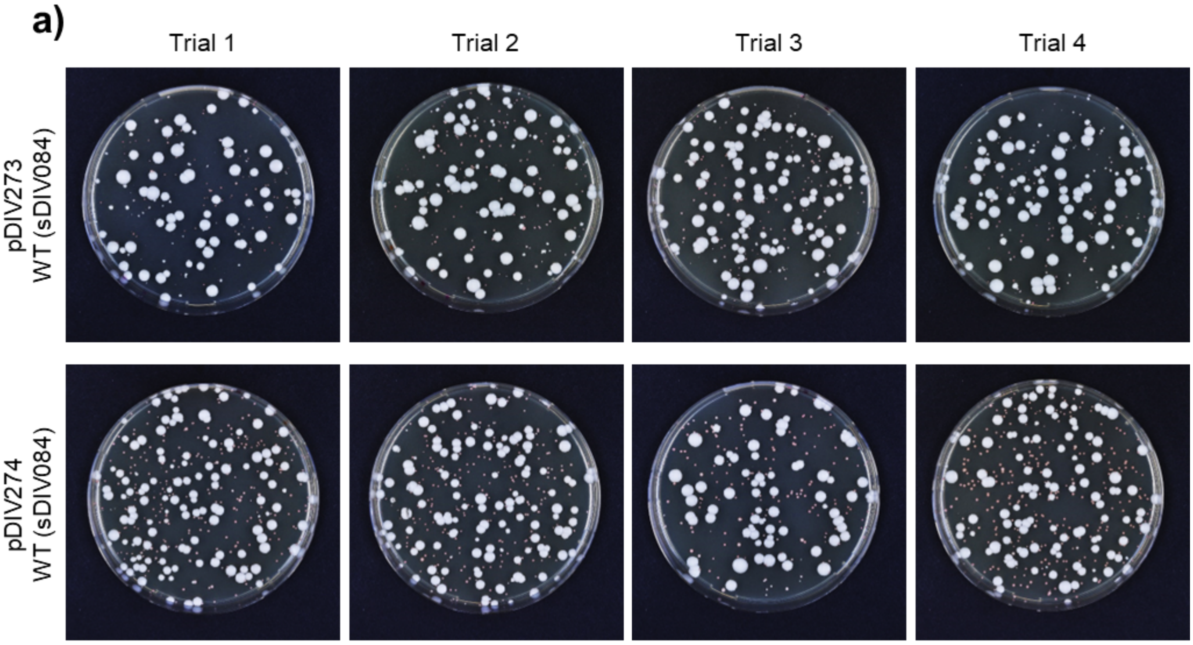


**Supplementary Figure S6.** Images of Transformation plates of *ADE2* experiment. Top row, four replicates transformed with pDIV273 plasmid, bottom row, with pDIV274 plasmid. Note, the significantly decreased fitness represented by significantly smaller size of red cells as compared to the white ones. Image contrast was enhanced for better color representation.

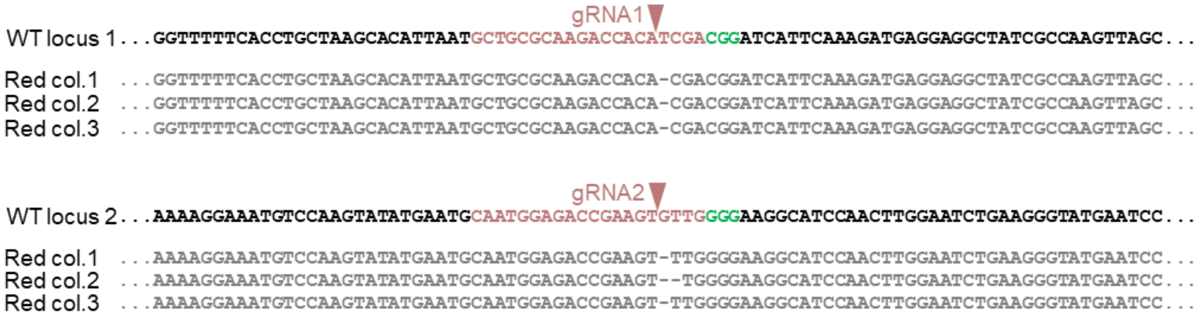


**Supplementary Figure S7.** An excerpt of Sanger sequencing results of ADE2 locus. Three red colonies resulting from transformants by pDIV273 (sgRNA1) plasmid (top panel) and from transformants by pDIV274 (sgRNA2) plasmid (bottom panel) were sequenced and aligned to the wild-type loci of *ADE2*. Light red text indicates 20 nt guiding sequences, green- PAM sequences.

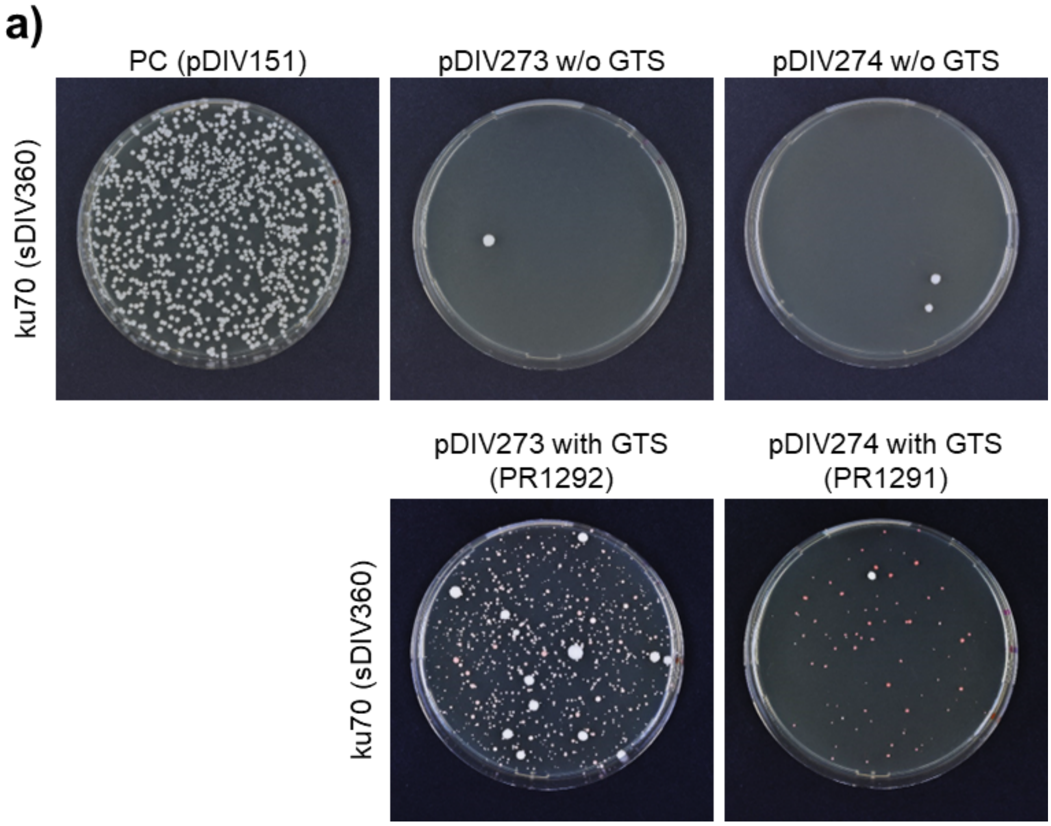


**Supplementary Figure S8.** Images of agar plates representing a TAPE experiment. Each plate represents equimolar amounts of plasmid DNA transformed. Note, only few colonies of *ku70* strain survive when sgRNA-Cas9 plasmids are transformed without repair templates (GTS).

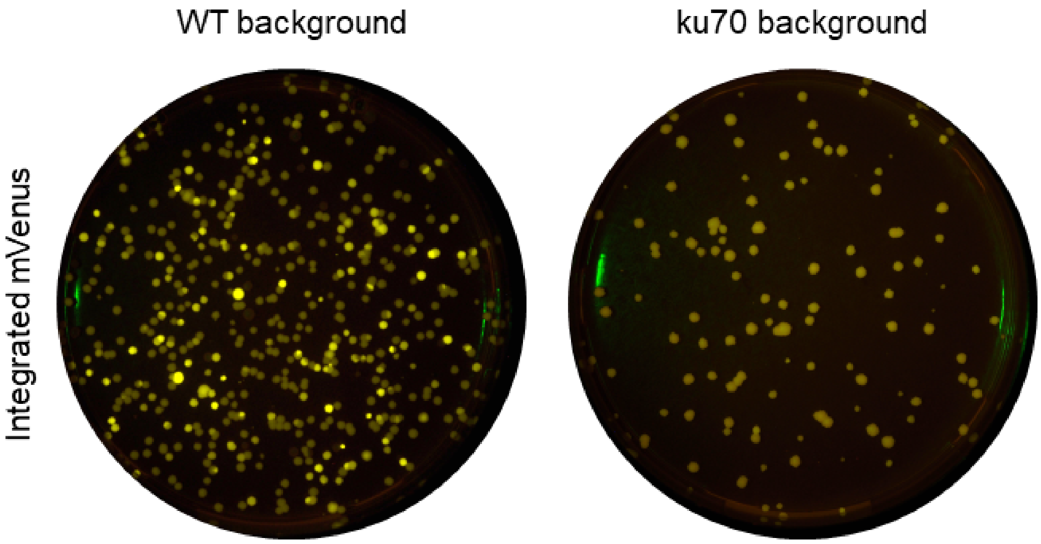


**Supplementary Figure S9.** Images of agar plates with yellow fluorescent colonies. The full plate pictures representing the snippets from Figure 2 in the main text. Left, a typical transformation (GTS-mVenus + sgRNA-Cas9 plasmid) plate image when mVenus expression cassette is transformed into WT (NHEJ proficient) strain. Right, a typical transformation (GTS-mVenus + sgRNA-Cas9 plasmid) plate image when mVenus expression cassette is transformed into ku70 (NHEJ deficient) strain.

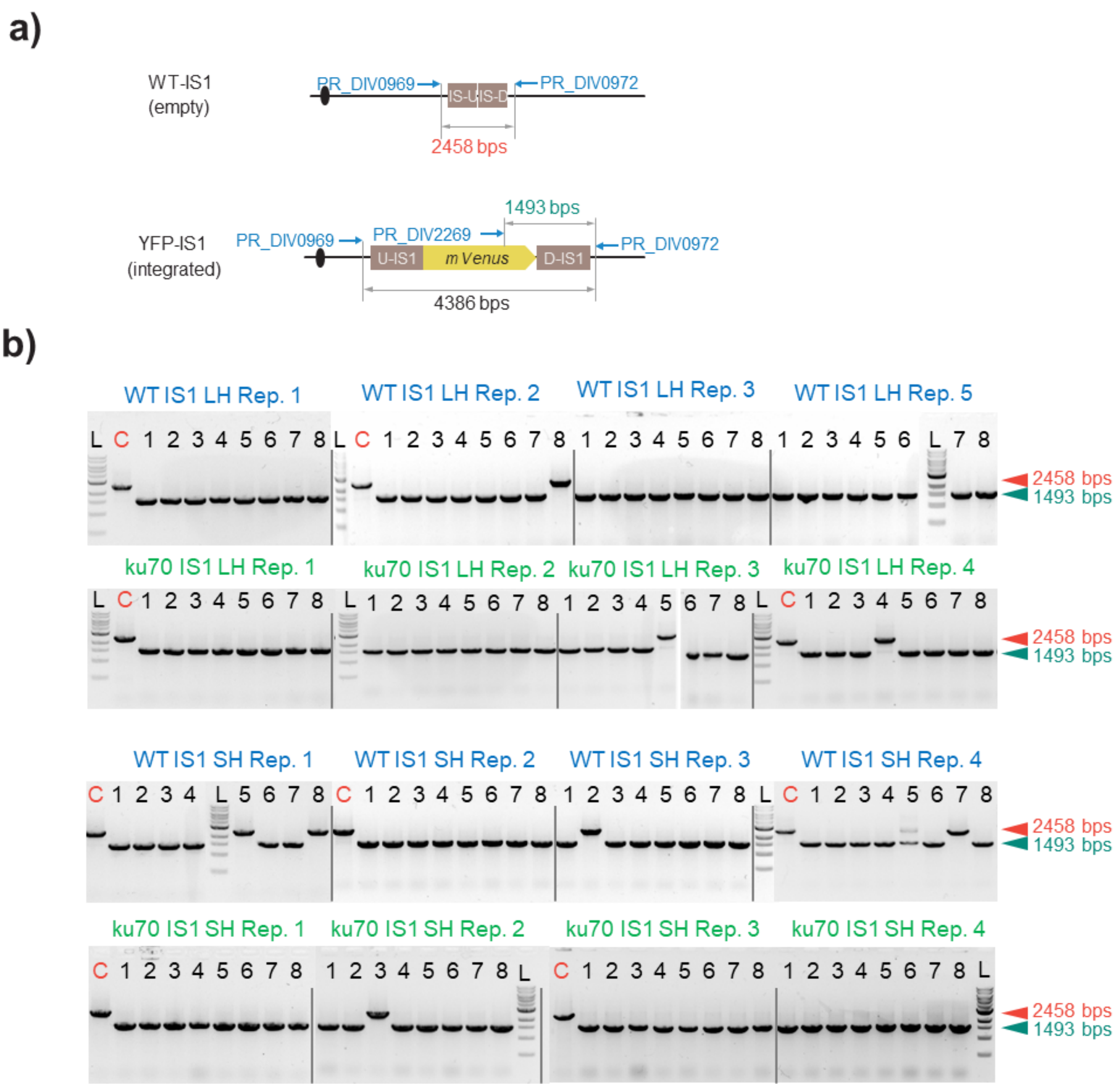


**Supplementary Figure S10.** Validation of integrated mVenus cassette into integration site (IS1). **a)** Schematic depiction of the IS1 locus, top wild-type (WT) and bottom, modified (YFP-IS1). Blue arrows indicate primer binding positions, and their names (blue text). Red text indicates expected band size if the locus is "empty" and green text - when mVenus is correctly integrated in the IS1 site. **b)** A composite image of agarose gels showing colony PCR results for IS1 locus. Blue text, bands obtained in NHEJ proficient (WT) background strain, and green text in NHEJ deficient (ku70) strain. Four biological replicates (Rep.) were tested in each strain background. L - 1 kb ladder (NEB), C - untransformed strain was used as a control.

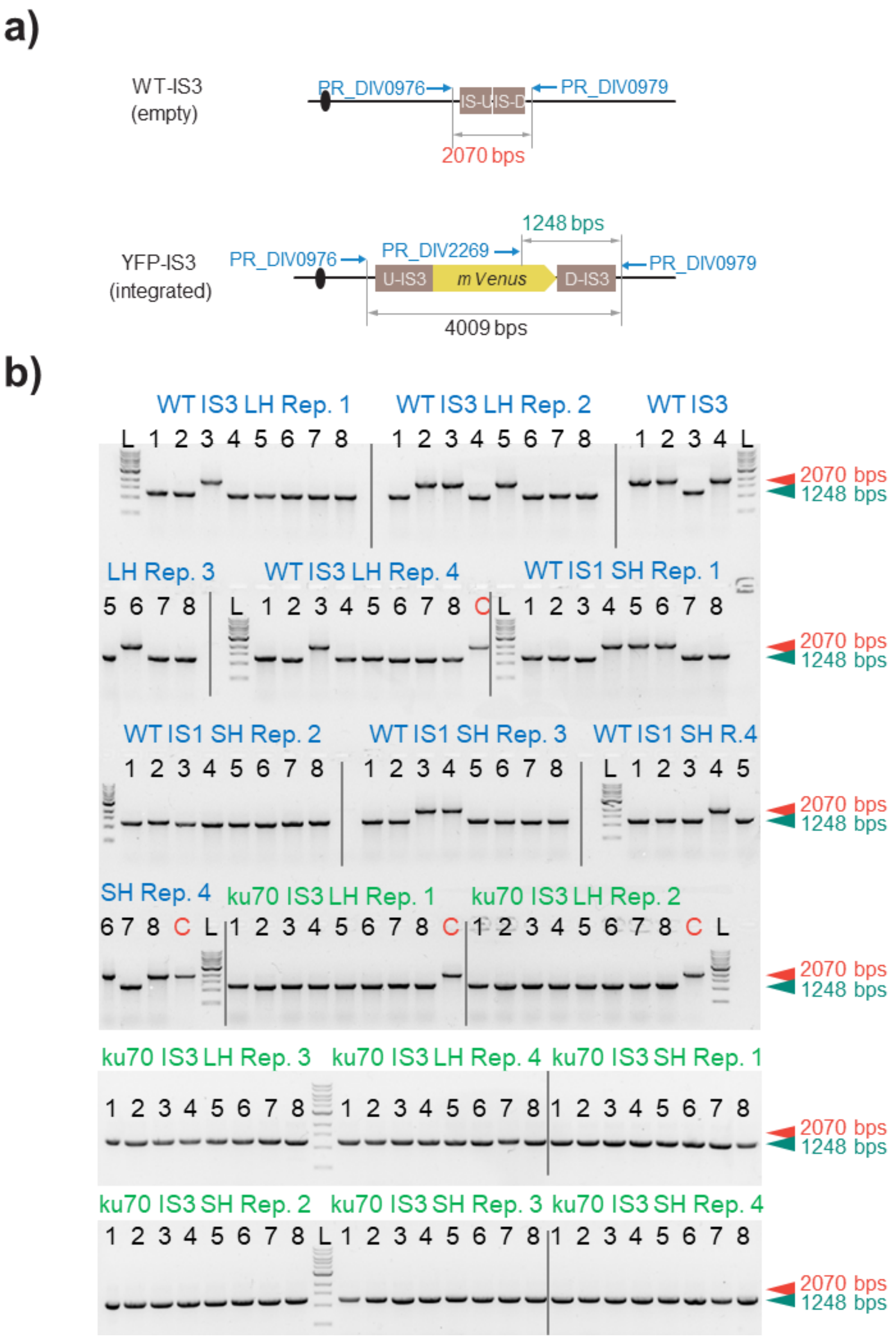


**Supplementary Figure S11.** Validation of integrated mVenus cassette into integration site (IS3). **a)** Schematic depiction of the IS3 locus, top wild-type (WT) and bottom, modified (YFP-IS3). Blue arrows indicate primer binding positions, and their names (blue text). Red text indicates expected band size if the locus is "empty" and green text - when mVenus is correctly integrated in the IS3 site. **b)** A composite image of agarose gels showing colony PCR results for IS3 locus. Blue text, bands obtained in NHEJ proficient (WT) background strain, and green text in NHEJ deficient (ku70) strain. Four biological replicates (Rep.) were tested in each strain background. L - 1 kb ladder (NEB), C - untransformed strain was used as a control.

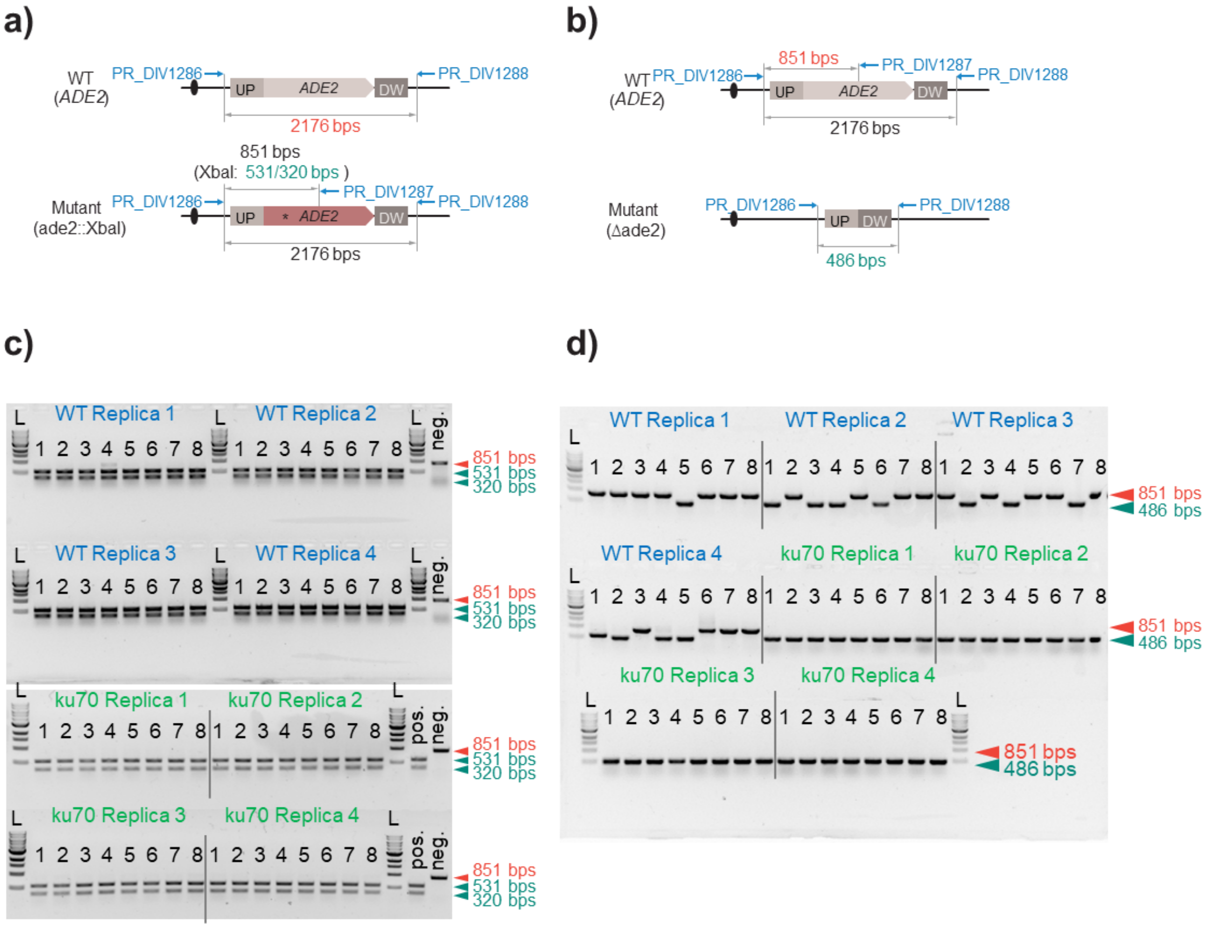


**Supplementary Figure S12.** Validation of modified ADE2 locus using short single stranded DNA oligonucleotides as GTS. **a)** Schematic depiction of the *ADE2* locus, top wild-type (WT) and bottom, modified (ade2::Xba**I**). Blue arrows indicate primer binding positions, and their names (blue text). Red text indicates expected band size if the locus is "*ADE2*". Green text shows sizes of two bands resulting after a PCR fragrant is digested with Xba**I** enzyme if ADE2 locus was correctly mutated. **b)** Schematic depiction of the *ADE2* locus, top wild-type (WT) and bottom, deleted (Δade2). Blue arrows indicate primer binding positions, and their names (blue text). Red text indicates expected band size if the locus is wild-type "*ADE2*". Green text shows band size if the *ADE2* gene was deleted. **c)** A composite image of agarose gels showing colony PCR results for *ADE2* locus mutation via ss-GTS. PCRs were treated with Xba**I** enzyme prior loading on gel. Blue text, bands obtained in NHEJ proficient (WT) background strain, and green text in NHEJ deficient (ku70) strain. Four biological replicates (Replica) were tested in each strain background. L - 1 kb ladder (NEB), pos. - positive control, neg- negative control. **d)** A composite image of agarose gels showing colony PCR results for *ADE2* locus deletion via ss-GTS. Blue text, bands obtained in NHEJ proficient (WT) background strain, and green text in NHEJ deficient (ku70) strain. Four biological replicates (Replica) were tested in each strain background. L - 1 kb ladder (NEB), pos. - positive control, neg- negative control.

**a)**

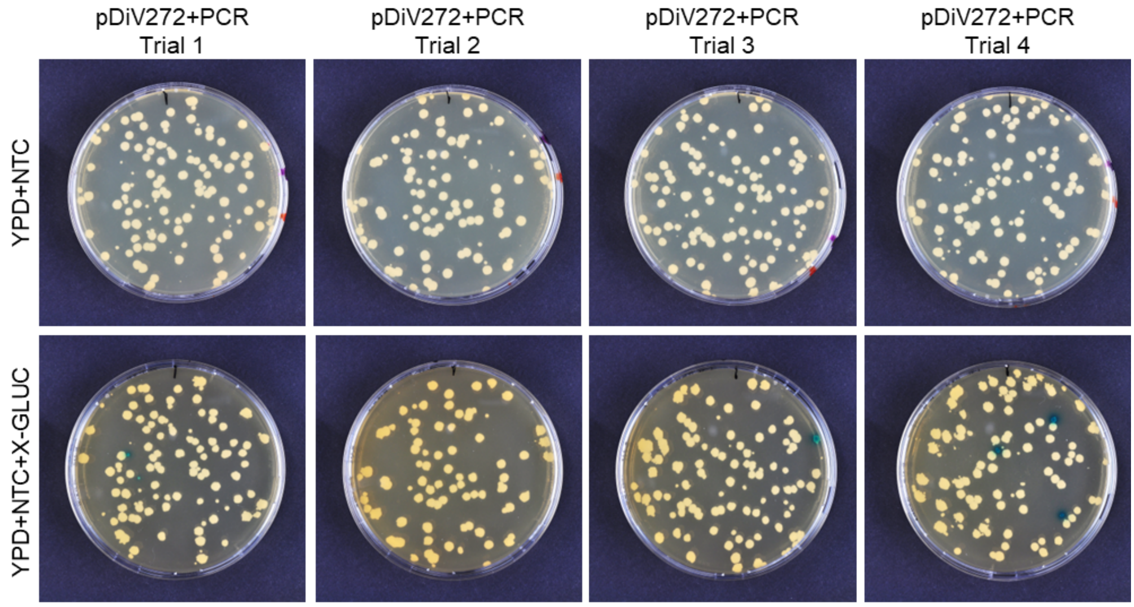


**b)**

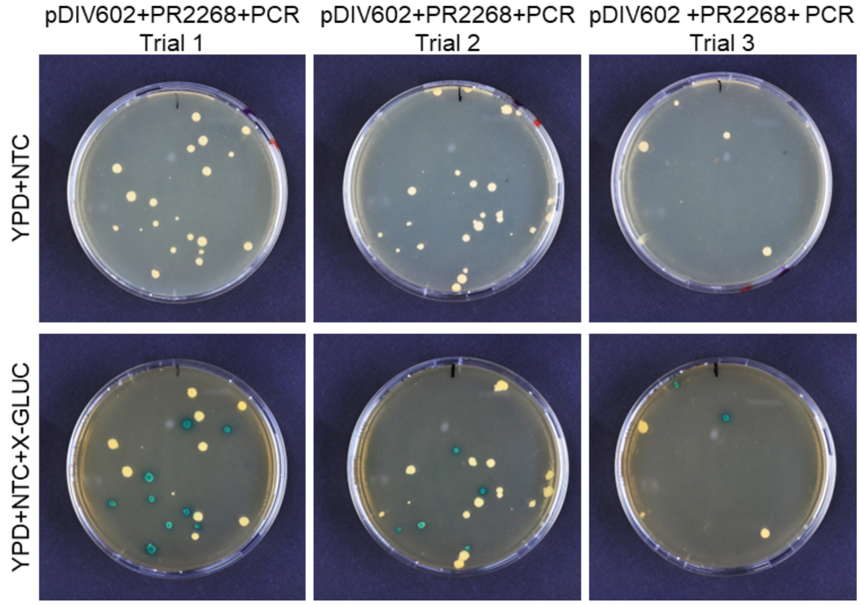


**Supplementary Figure S13.** Images of agar plates of *ku70* reversion and *PEP4* deletion experiment. **a)** Transformation plates (top row) and corresponding replica plates (bottom row) of the *ku70* reversion. **b)** Transformation plates (top row) and corresponding replica plates (bottom row) of the multiplex experiment for *PEP4* deletion and *ku70* reversion

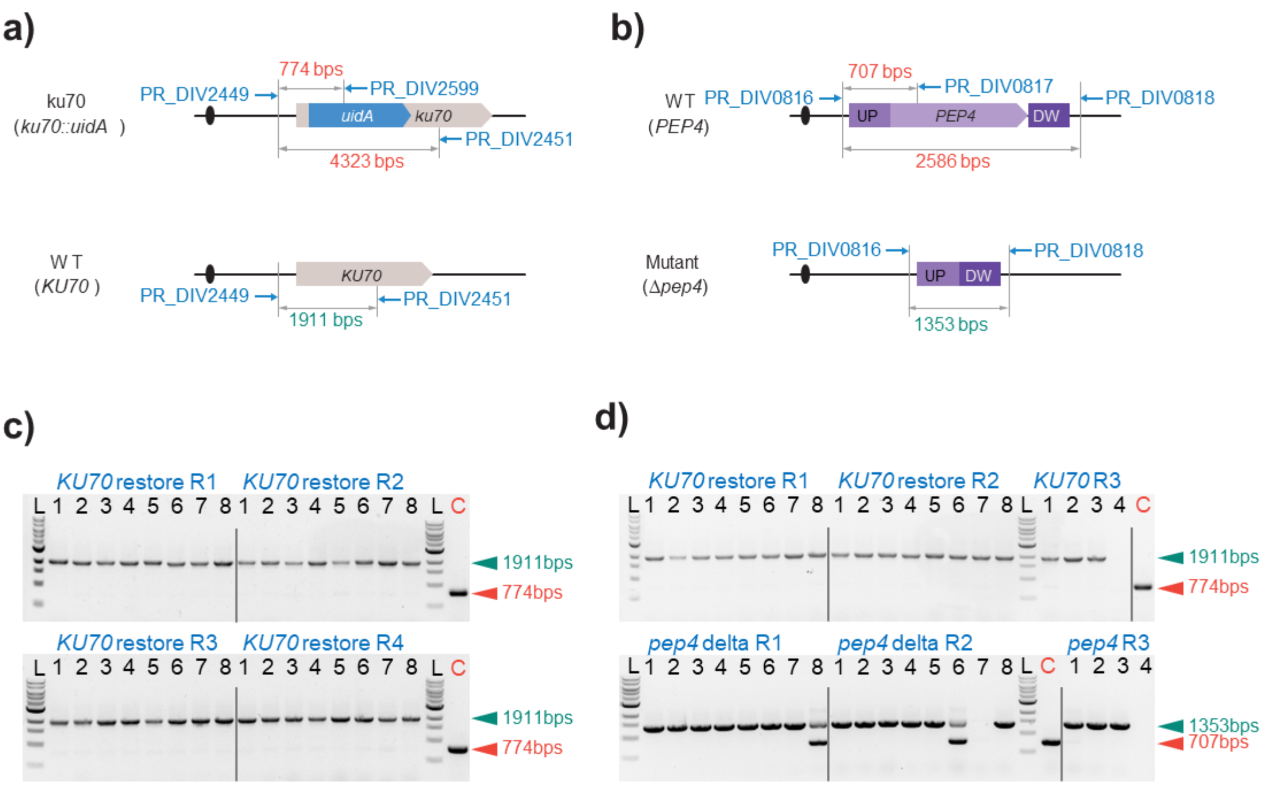


**Supplementary Figure S14.** Validation of modified *KU70* locus *PEP4* gene deletion. **a)** Schematic depiction of the *KU70* locus, top modified (*ku70::uidA*) and bottom, wild-type (*KU70*). Blue arrows indicate primer binding positions, and their names (blue text). Red text indicates expected band size if the locus is “*ku70::uidA*”. Green text shows the size of a PCR band when KU70 was restored. **b)** Schematic depiction of the *PEP4* locus, top wild-type (WT) and bottom, deleted (Δpep4). Blue arrows indicate primer binding positions, and their names (blue text). Red text indicates expected band size if the locus is wild-type “*PEP4*”. Green text shows band size if the *PEP4* gene was deleted. **c)** A composite image of agarose gels showing colony PCR results for *KU70* locus. Four biological replicates (R#) were tested in each strain background. L - 1 kb ladder (NEB), C - untransformed sDIV291 strain was used as a control. **d)** A composite image of agarose gels showing colony PCR results for simultaneous recovery of *KU70* (top Gel) and *PEP4* (bottom Gel) gene deletion via ss-GTS. Three biological replicates (R#) were tested in each strain background. L - 1 kb ladder (NEB), C - untransformed sDIV291 strain was used as a control.
